# Holistic monitoring of freshwater and terrestrial vertebrates by camera trapping and environmental DNA

**DOI:** 10.1101/2022.11.23.517571

**Authors:** Anne Marie Rubæk Holm, Steen Wilhelm Knudsen, Malene Månsson, Ditte Elmgreen Pedersen, Pauli Holm Nordfoss, Daniel Klingberg Johansson, Marthe Gramsbergen, Rasmus Worsøe Havmøller, Eva Egelyng Sigsgaard, Philip Francis Thomsen, Morten Tange Olsen, Peter Rask Møller

## Abstract

The anthropogenic impact on the world’s ecosystems is severe and the need for non-invasive, cost-effective tools for monitoring and understanding those impacts are therefore urgent. Here we combine two such methods in a comprehensive multi-year study; camera trapping (CT) and analysis of environmental DNA (eDNA), in river marginal zones of a temperate, wetland Nature Park in Denmark. CT was performed from 2015 to 2019 for a total of 8,778 camera trap days and yielded 24,376 animal observations. The CT observations covered 87 taxa, of which 78 were identified to species level, and 73 were wild native species. For eDNA metabarcoding, a total of 114 freshwater samples were collected from eight sites in all four seasons from 2017 to 2018. The eDNA results yielded a total detection of 80 taxa, of which 74 were identified to species level, and 65 were wild native species. While the number of taxa detected with the two methods were comparable, the species overlap was only 20 %. In combination, CT and eDNA monitoring thus yielded a total of 115 wild species (20 fishes, four amphibians, one snake, 23 mammals and 67 birds), representing half of the species found via conventional surveys over the last ca. 20 years (83% of fishes, 68 % of mammals, 67 % of amphibians, 41 % of birds and 20 % of reptiles). Our study demonstrates that a holistic approach combining two non-invasive methods, CT and eDNA metabarcoding, has great potential as a cost-effective biomonitoring tool for vertebrates.

## 1 Introduction

Freshwater ecosystems and their bordering terrestrial habitats cover a small fraction of the Earth’s surface yet support about a third of all known vertebrate species (Strayer & Dudgeon, 2010). These habitats are highly vulnerable to human activities, such as urban development, agriculture, nutrient and waste-water runoff, aquaculture, fisheries and damming (Arthington et al., 2006; Dudgeon et al., 2006; Naiman et al., 2002), necessitating efficient methods for monitoring their biodiversity. Conventional methods for such monitoring include direct visual or acoustic observations, or indirect detections via e.g., tracks, scat or sloughed feathers or fur. In the past decades, camera trapping (CT) has proven to be a minimally invasive and highly efficient method for detection and long-term monitoring of vertebrate biodiversity (e.g., Ahumada et al., 2013; Mugerwa et al., 2013; Silveira et al., 2003). The method allows detection of elusive (Trolle & Kéry, 2005), rare (Azlan and Lading, 2006) and novel species (Rovero et al., 2008), and while CTs are often used to study mammals in tropical areas (Burton et al., 2015; Havmøller et al., 2019), they have also proven effective in temperate forests and open areas (Rovero et al., 2014; Parsons et al., 2018). More recently, environmental DNA (eDNA) analysis has emerged as another cost-effective and non-invasive method for biodiversity monitoring (Ficetola et al., 2008; Taberlet et al., 2012; Thomsen & Willerslev, 2015). This method has been used for species inventories across a wide range of habitat types, although most applications to date are in aquatic systems (e.g., Pedersen et al., 2015; Goldberg et al., 2016; Thomsen et al., 2012).

All biomonitoring methods have their strengths and weaknesses in terms of taxonomic coverage, ease of use, survey effort and requirements of taxonomic expertise, and not one method can capture the entire vertebrate diversity of an ecosystem. For instance, combining eDNA metabarcoding and CTs for monitoring of marine fishes has resulted in detection of a larger richness than any of these approaches alone (Stat et al., 2018; Boussarie et al., 2018). Similarly, metabarcoding analysis of eDNA from stream water (Lyet et al., 2021) and terrestrial sediments (Leempoel et al., 2020) combined with CTs has been found to be efficient for monitoring terrestrial mammals. The number of such vertebrate studies combining water eDNA and CTs is growing rapidly, in covering all sorts of habitats from reefs (Stat et al., 2018; Boussarie et al., 2018) to ponds (Mas-Carrió et al., 2022; Harper et al., 2019).

Here, we combine one year of aquatic eDNA sampling and four years of CT data collection to investigate the vertebrate fauna in a Danish wetland and Nature Park in temperate Northern Europe. We provide an updated inventory of the diversity of species in the park, their commonness and conservation status, and evaluate the complementarity, strengths, and weaknesses of monitoring aquatic eDNA versus monitoring with CTs and compare our results with baseline data for the same locality collected by conventional biodiversity monitoring methods over the past two decades.

## 2 Materials and methods

### 2.1 Study site

Field work was performed at Nature Park Åmosen (hereafter referred to as Åmosen), West Zealand, Denmark (N 55.618860, W 11.329161). Åmosen comprises a stream system of approximately 45 km from Undløse in the east to the Great Belt in the west (Figure 1). It consists of a mixed set of habitats including streams, wetlands, forests, fens, meadows, bogs, and thickets, as well as agriculture and some urban development. Åmosen holds a unique flora and fauna including several red-listed species and about 80% of the park is designated as a Natura 2000 area (area no. 156, H137 and area no. 157, H138, F100) (Schmidt, 2017; Naturstyrelsen, 2016a; Naturstyrelsen, 2016b).

**FIGURE 1.**
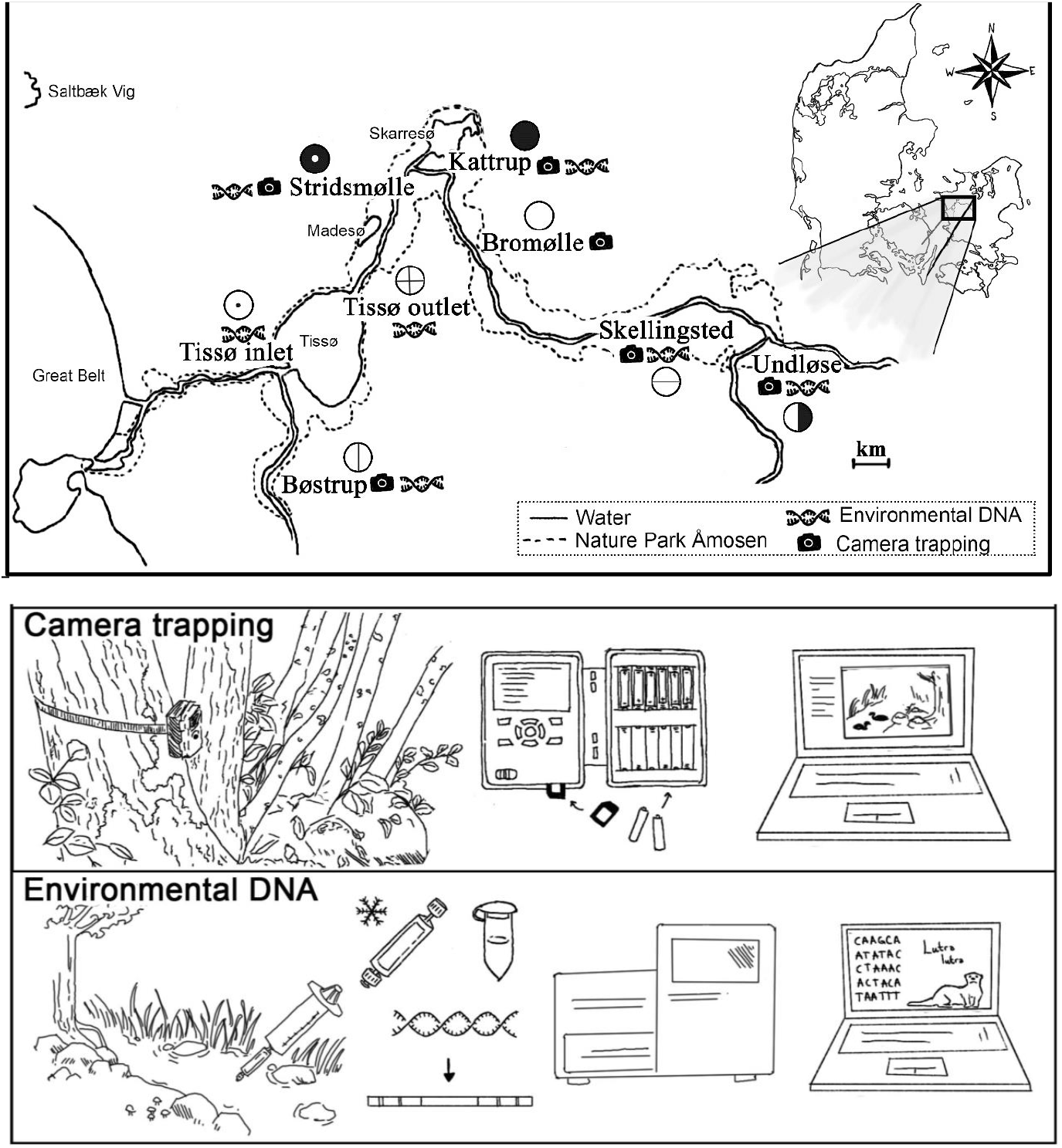
The Åmosen Nature Park sampling sites as well as schematic illustration of the camera trapping and environmental DNA methods used to monitor vertebrate diversity. Illustrations by AMRH.

### 2.2 Camera trapping

We monitored the vertebrate fauna of Åmosen by deploying up to 16 camera traps (CTs) at six locations over a period of four years from the 20^th^ of May 2015 to the 12^th^ of August 2019 (Table 1). The number of CTs varied by location and season, as some sites were more suitable for deployment than others, and as cameras were occasionally lost due to theft and flooding. We used a water-resistant CT model (IR PLUS BF HD) equipped with a passive infrared sensor and a 940 nm light-emitting diode flash source. All CTs were placed facing the catchment area and angled to cover both the stream and the opposite stream bank, as suggested by Matsubayashi et al., (2006). The CTs were programmed to record photos and/or 10 second videos with normal sensitivity and no trigger interval, and no bait or lures were used. Batteries and memory cards were replaced at regular intervals.

**Table 1.** Sampling details for each study site (see Figure 1) and overall. Number (N) of camera traps (CTs) at each study site, camera days (CDs) (collected from 20th of May 2015 to 12th of August 2019), and number of camera events (CE’s) (observations of animal with at least a 30-minute interval). eDNA samples were collected all through 2018 (January*, February, March, May, September, and December), a few samples were taken in the end of 2017 (September** and October**). *only for Stridsmølle. **only for Kattrup and Stridsmølle.

|  |  | Bromølle | Bøstrup | Katstrup | Skellingsted | Stridsmølle | Tissø inlet | Tissø outlet | Undløse | All sites |
| --- | --- | --- | --- | --- | --- | --- | --- | --- | --- | --- |
|  |  | Stream, forest and settlements | Stream, reed beds and agriculture | Stream in forested landscape | Regulated stream in agricultural landscape | Stream in forested landscape | Lake in agricultural landscape | Lake in agricultural landscape | Regulated stream in agricultural landscape |  |
| **Site characteristics** |  |  |  |  |  |  |  |  |  |  |
| Camera | Period | 6/4-2017-12/8-2019 | 6/4-2017-12/8-2019 | 20/5-2015-12/8-2019 | 6/4-2017-12/8-2019 | 20/5-2015-12/8-2019 | - | - | 6/4-2017-12/8-2019 |  |
|  | N | 4 | 1 | 8 | 2 | 1 | - | - | 2 | 18 |
|  | CDs | 670 | 97 | 6,487 | 514 | 1,007 | - | - | 106 | 8881 |
|  | CEs | 770 | 107 | 6907 | 546 | 578 | - | - | 181 | 9089 |
|  | Taxa | 34 | 20 | 71 | 37 | 34 | - | - | 34 | 87 |
| eDNA | Period | - | 27/2-13/12-2018 | 27/9-2017-13/12-2018 | 27/2-13/12-2018 | 17/9-2017-13/12-2018 | 27/2-13/12-2018 | 27/2-13/12-2018 | 27/2-13/12-2018 |  |
|  | N | - | 15 | 19 | 15 | 22 | 14 | 14 | 15 | 114 |
|  | Taxa | - | 40 | 50 | 46 | 49 | 45 | 46 | 42 | 50 |
| Taxa | Aves | 20 | 22 | 53 | 23 | 26 | 21 | 23 | 26 | 74 |
|  | Mammalia | 14 | 9 | 24 | 20 | 17 | 7 | 5 | 15 | 29 |
|  | Actinopterygii | 0 | 14 | 19 | 18 | 17 | 16 | 17 | 12 | 20 |
|  | Amphibia | 0 | 2 | 1 | 1 | 3 | 1 | 1 | 3 | 4 |
|  | Squamata | 0 | 0 | 0 | 1 | 0 | 0 | 0 | 0 | 1 |
|  | Domestic | 2 | 4 | 7 | 4 | 4 | 4 | 3 | 4 | 9 |
|  | Total | 34 | 47 | 97 | 63 | 63 | 45 | 46 | 56 | 137 |

Photos and videos from CTs were manually examined and identified to the lowest possible taxonomic level based on morphological traits, movement patterns and sounds with help from taxonomic experts at the Natural History Museum of Denmark. To avoid artificial inflation of observations, a camera event (CE) was defined as all detections of a certain species within 30 minutes at the same location (O’Brien et al., 2003; Zimmermann & Rovero, 2016). To assess the commonness of each taxon, we estimated the relative abundance index (RAI) as the number of CEs of a given taxon per 100 camera trap days (O’Brien, 2011; Rovero et al., 2014), and the naïve occupancy (NO) as the proportion of sites that recorded at least one CE of the target species (e.g., Jenks et al., 2011; Rovero et al., 2014; Hedwig et al., 2017).

### 2.3 Environmental DNA

In addition to monitoring by CTs, we performed eDNA-based monitoring of vertebrates by collection of water samples from September 2017 to December 2018 (Table 1, Figure 1). At each sampling event, two to three sample replicates were collected. Each sample replicate consisted of up to 500 ml of water taken with a 60 mL syringe (Soft-Ject, HSW, Tuttlingen, Germany) and filtered through a Sterivex filter unit of 0.22 μm pore size (polyethersulfone, Merck Millipore, Germany). The samples were transported in a cooler and stored at −18 °C until DNA extraction.

All laboratory work was performed in separate laboratories designated for DNA extraction, pre-PCR, and post-PCR procedures, respectively. Environmental DNA was extracted from the filters using the DNeasy Blood & Tissue Kit (Qiagen GmbH, Hilsen, Germany) with a modified protocol (Sigsgaard et al., 2020; Spens et al, 2017). Polymerase chain reaction (PCR) amplification was performed using the primer set Mamm01 (mamm01_F:5’-CCGCCCGTCACCCTCCT-3’, mamm01_R: 5’-GTAYRCTTACCWTGTTACGAC-3’) (Taberlet et al., 2018), and the primer set MiFish-U (MiFish-U_F: 5’-GTCGGTAAAACTCGTGCCAGC-3’, MiFish-U_R: 5’-CATAGTGGGGTATCTAATCCCAGTTTG-3’) (Miya et al., 2015). These primer sets target regions of approximately 59 bp and 170 bp (excluding primers), respectively, around 390-400 bp apart in the 12S mitochondrial gene. The DNA extracts of the three sample replicates from each site were pooled into one sample. Subsequent analyses were done on the resulting 40 sample pools. Setup of quantitative PCR (qPCR) and PCR for meta barcoding with reagents, volumes, concentrations, and thermocycler conditions are provided in the supplementary material (supporting text A, B and tables S1-S10). Each PCR setup included one PCR replicate of each eDNA sample pool, negative PCR controls and a positive mock sample comprising genomic DNA from 24 exotic species unlikely to be found in Denmark, including mammals, fish, and a frog (Olds et al., 2016; Thomsen et al., 2016), but we failed to include extraction controls. Six PCR replicates were run of each sample pool, for each primer set, giving a total of 12 libraries (supplementary material table S1-S8).

All PCR products were verified on a 2% agarose gel stained with GelRed (Biotium). From each of the 12 libraries (2*3 replicates for MiFish-U and 2*3 replicates for Mamm01) (supplementary material table S1-S8) we pooled 10 μL to a total of 120 μL. The 120 μL was then purified using the MinElute (Qiagen) PCR purification kit (cat. no. 28006), following the supplied protocol with modifications (supplementary material, supporting text C). Twelve 150 bp paired-end libraries (six for the Mamm01 primer set and six for the MiFish-U primer set) were prepared with an Illumina TruSeq DNA PCR-free LT Sample Prep kit (Illumina, San Diego, California), spiked with 8% phiX, and sequenced on two Illumina MiSeq3 flow cells (six libraries on each, the Mamm01 libraries on one flow cell, and the MiFish on another flow cell) at the GeoGenetics Sequencing Core, University of Copenhagen, Denmark.

Sequence reads were demultiplexed using the software package Cutadapt (Martin, 2011) and a custom python script (available at https://github.com/tobiasgf/Bioinformatic-tools/tree/master/Eva_Sigsgaard_2018) (Sigsgaard et al., 2020). Reads shorter than 10 bp or including ambiguities or with >2 expected errors were removed (Sigsgaard et al., 2020). We then used DADA2 (Callahan et al., 2016) to correct PCR and sequencing errors in the raw sequencing output, and forward and reverse reads with a minimum of 5 bp overlap and no mismatches were then merged. Sequences were blasted against the National Center for Biotechnology Information (NCBI) GenBank database using BLASTn (Altschul et al., 1990) on the 20^th^ of March 2020. BLASTn settings were set to a maximum of 3000 hits per query (-max_target_seqs 3000), minimum thresholds of 90 % query coverage per high-scoring segment pair (-qcov_hsp_perc 90), and 80 % sequence similarity (- perc_identity 80). The output format was set to: -outfmt “6 std qlen qcovs sgi sseq ssciname staxid”. BLAST hits displaying incomplete final query coverage were removed. We then classified hits taxonomically in R v.3.6 (R Core team, 2020), using the package ‘taxize’ (Chamberlain and Szocs, 2013). To reduce data processing time, BLAST hits were then compared against a list of regional vertebrate species and hits to species that are exotic to northern Europe were removed (Figure 1). We removed exotic species with >95% match match with the mock species (supplementary material Table S10). The naïve occupancy (NO) was calculated across all eDNA samples and for each study site, respectively, as the number of eDNA sites/samples where a given taxon was detected divided by the total number of eDNA sites/samples (Table 1).

### 2.4 Method comparison

To compare our CT- and eDNA-based species detections with previous biodiversity monitoring efforts, we summarized data from conventional vertebrate surveys performed in Åmosen over the last two decades (2000-2020). Species presence data was compiled from BirdLife Denmark (DOF) (Grell, 1998 and recent data from Michael Fink), Baagøe and Jensen (2007), Carl and Møller (2012), and the Danish species portal Arter.dk, as well as from additional direct visual observations, trapping, excrements, tracks, roadkill done during the CT and eDNA field work and museum collections.

## 3 Results

### 3.1 Camera trapping

The camera trapping yielded a total of 8,778 camera days with 24,376 animal sightings across 8,674 CEs. These sightings represented 87 vertebrate taxa, of which 78 (90%) were identified to species level (Table 1). While birds (57 taxa) and mammals (29 taxa) dominateda grass snake (*Natrix natrix*) and a northern pike (*Esox lucius*) caught by a grey heron (*Ardea cinerea*) were also observed (Figure 2, 3, Supplementary Table S9). Most observed taxa were wild species, but domestic animals such as cat (*Felis catus*), dog (*Canis lupus*), cattle (*Bos taurus*), chicken (*Gallus gallus*), and Muscovy duck (*Cairina moscata*) were also detected. The taxa differed markedly in detection frequency with 53% of the taxa being detected in less than 10 CEs and only 18% of the taxa being observed at more than 100 CEs (Figure 2a; Supplementary Table S9). The most observed bird was the mallard (*Anas platyrhynchos*) with a total of 1,422 CEs, amounting to an RAI of 16.2 (detection at 16.2% of all CEs on average) and an NO index of 1.0 (detection at all monitoring sites) (Figure 2b-c). Other frequently and/or widely detected birds included common wood pigeon (*Columba palumbus*), grey heron and Eurasian blackbird (*Turdus merula*). The mammal accounting for the most CEs was the roe deer (*Capreolus capreolus*), although this species was only observed at half of the sites (CEs=1172; RAI=13.4; NO=0.50), whereas the brown rat (*Rattus norvegicus*) was both frequently and widely observed (CEs=719; RAI=8.2; NO=1.0). Other frequently and/or widely encountered taxa included pine marten (*Martes martes*) and other mustelids, red squirrel (*Sciurus vulgaris*) and red fox (*Vulpes vulpes*). A rare surprise was the Eurasian otter (*Lutra lutra*), thought to be locally extinct at the time of the study but found here in 35 CEs, and first time May 28, 2016 (Figure 3). Human activity was recorded at all six study sites, in a total of 472 sightings (472/24,376=2%) and 111 individual CEs (111/8,674=1%).

**FIGURE 2.**
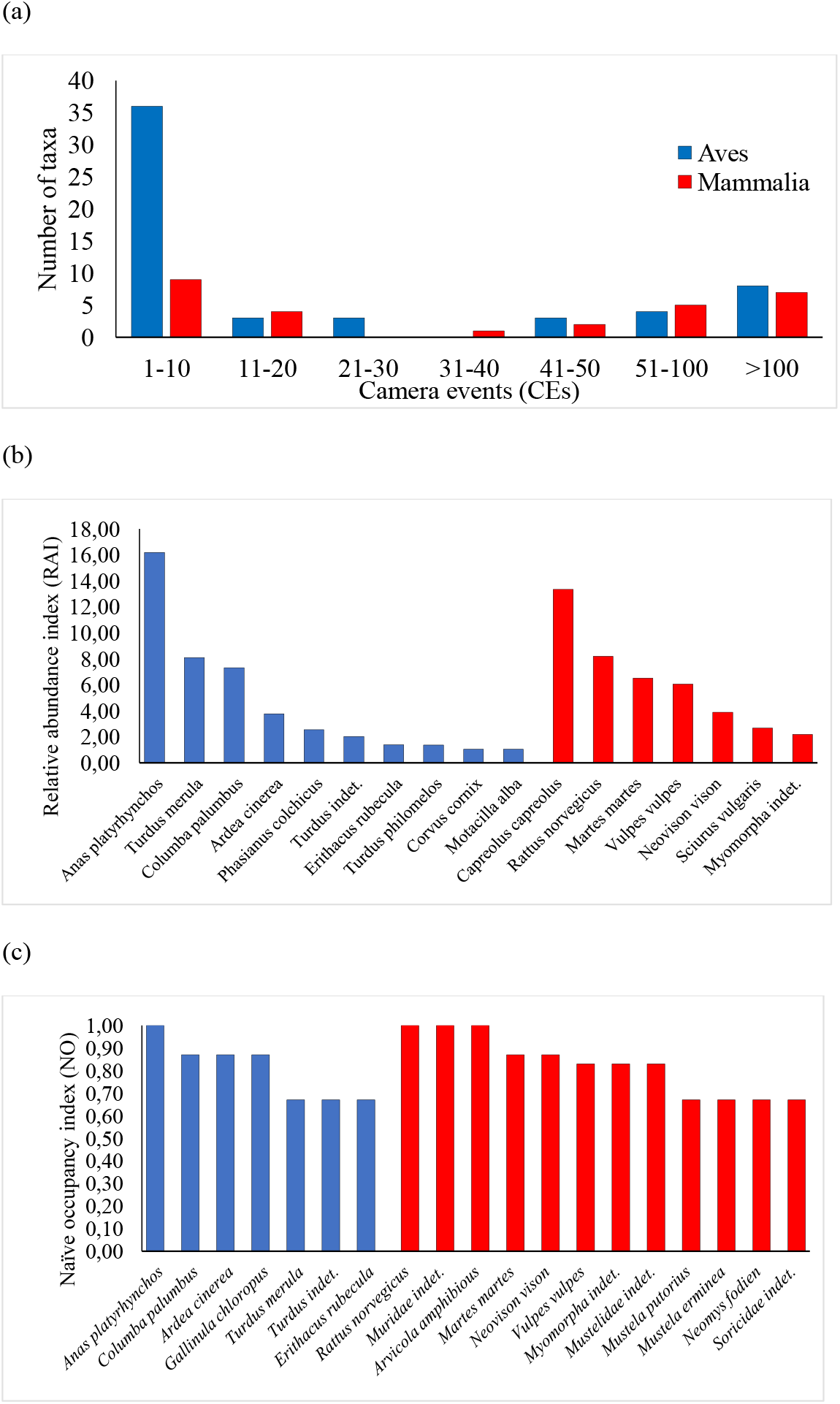
The bird and mammal taxa detected by camera traps differed greatly in their abundance and occupancy. (a) The number of taxa in different CE categories with a few very common taxa and many rare. (b) The most abundant bird and mammal taxa defined by a relative abundance index RAI>1.0. (c) The most common taxa defined by naïve occupancy index. A full species list is provided in Supplementary Table S9.

**FIGURE 3.**
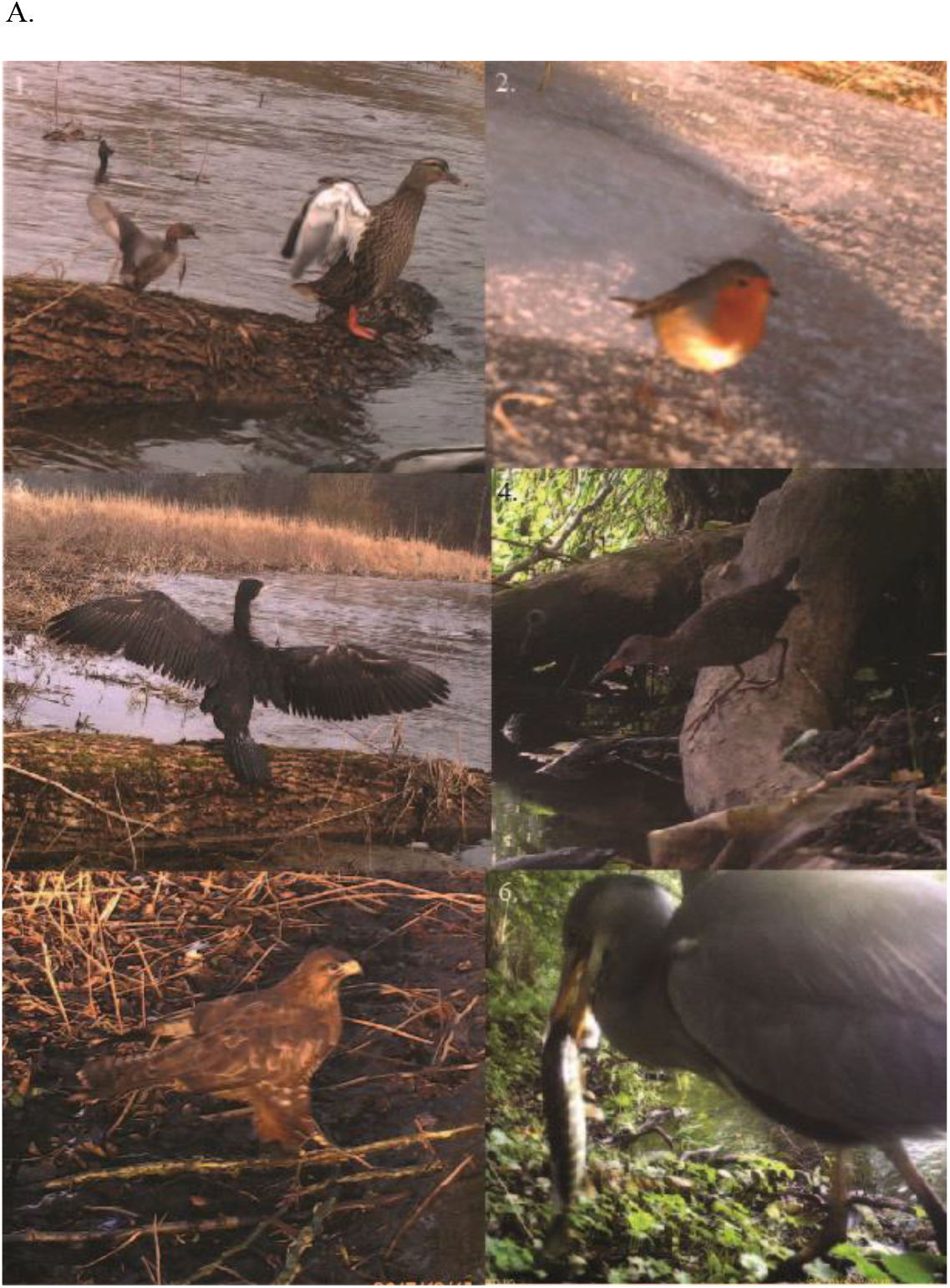
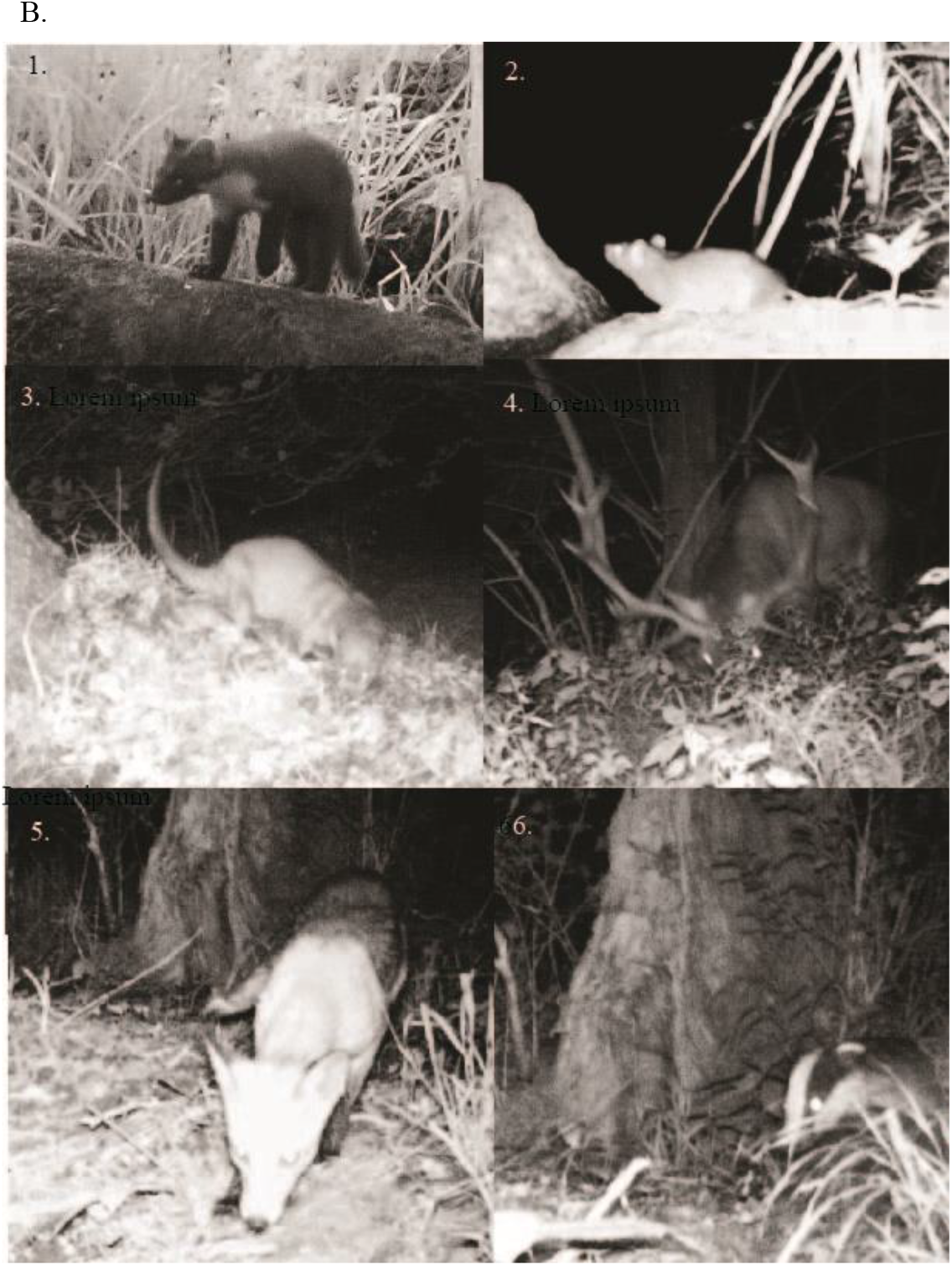
Examples of animals observed by CTs in Åmosen. (a) birds and fish 1. *Tachybaptus ruficollis* and *Anas platyrhynchos*, 2. *Erithacus rubecula*, 3. *Phalacrocorax carbo*, 4. *Rallus aquaticus*, 5. *Buteo buteo*, 6. *Ardea cinerea* and *Esox lucius*, and (b) mammals (1. *Martes martes*, 2. *Rattus rattus*, 3. *Lutra lutra*, 4. *Cervus elaphus*, 5. *Vulpes vulpes* and 6. *Meles meles.*

### 3.2 Environmental DNA

The Illumina MiSeq platform produced a total of 25,408,796 raw paired-end reads. After removing mock sample species, non-target species (e.g., prokaryotes and fungi) and human reads, a total of 12,154,093 reads from target vertebrates remained, which 48% of the reads being retained. Across the two primer sets, in the proportion of reads retained, matching vertebrates, 4% were identified as non-target vertebrates: The proportion of non-vertebrate sequence reads was much higher for the Mamm01 primer set (50-65%) than for the MiFish-U primer set (10-20 %). The retained sequence reads represented 80 taxa, of which 74 were identified to species level (Table 1, Supplementary Table S9). Both primer sets amplified mitochondrial DNA (mtDNA) from amphibians, fish and mammals, but while 49 taxa were identified by both primer sets, nine taxa were solely identified by the Mamm01 primer set and 22 taxa solely by the MiFish-U primer set. As expected, the MiFish-U primer set yielded more fish species than the Mamm01 primer set, but the Mamm01 primer set did not yield more mammal species (Supplementary Figure 1).

Humans were detected at all study sites, and nine of the detected species were domestic, including cat, cattle, chicken, dog, horse (*Equus ferus*), Muscovy duck, pig (*Sus scrofa*), sheep (*Ovis aries*), and turkey (*Meleagris gallopavo*). The common roach (*Rutilus rutilus*), and two other taxa dominated the eDNA data with more than one million sequence reads per taxa, while 20 of the taxa were detected in less than 1000 reads and five taxa in less than 100 reads (Figure 4a). The NO analysis also revealed large differences in species occupancy with a few bird, (domestic) mammals and fish taxa being detected at all eDNA study sites, including undetermined ducks, mallard (*Anser platyrhynchos*), Eurasian coot (*Fulica atra*), cow, pig, dog, undetermined arvicolines (voles and muskrats), common roach, Eurasian perch (*Perca fluviatilis*), ide (*Leuciscus idus*), northern pike, European eel (*Anguilla anguilla*) and rudd (*Scardinius erythrophthalmus*) (Figure 4b; Supplementary Table S9).

**FIGURE 4.**
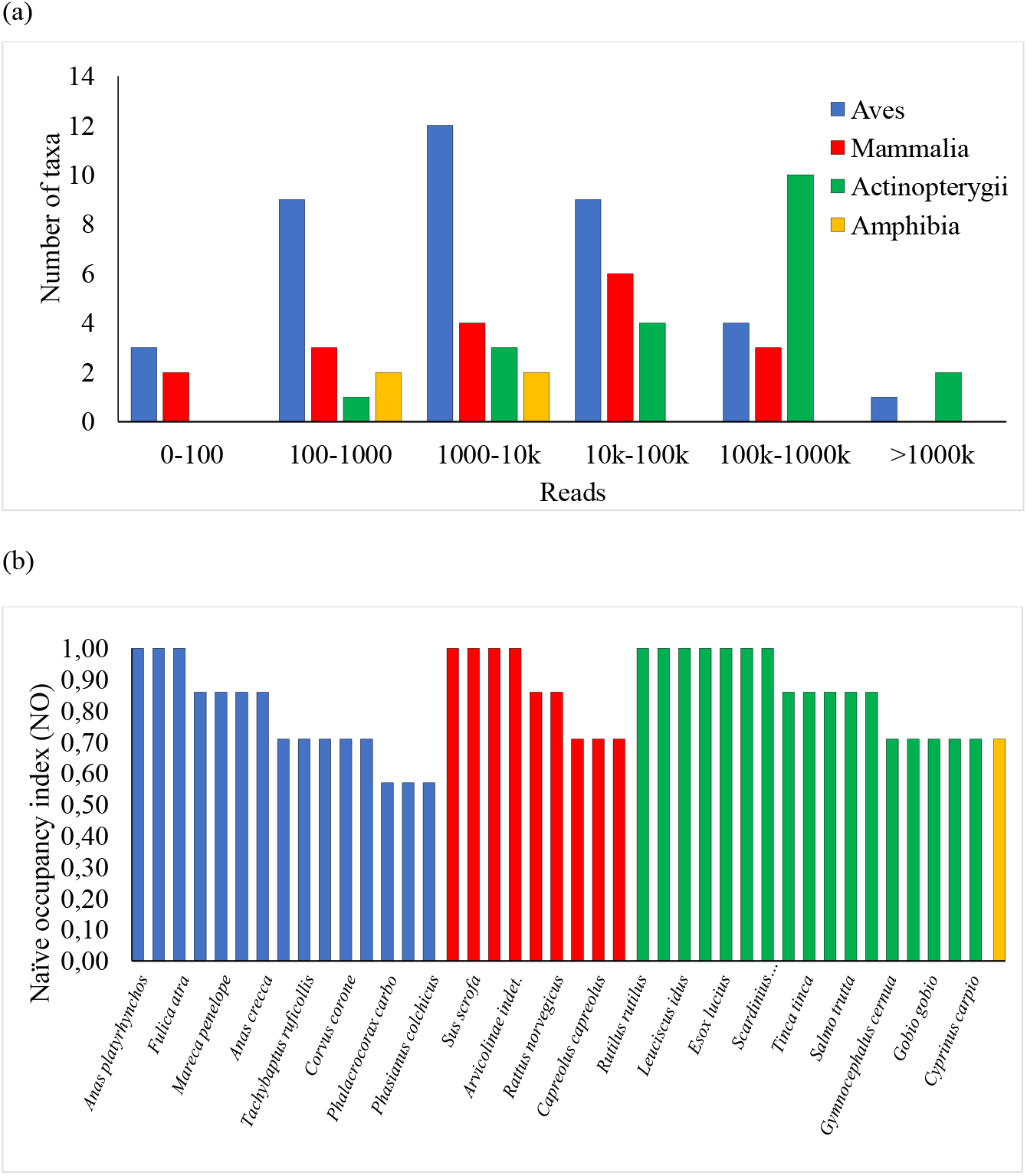
The bird, mammal, fish and amphibian taxa detected by eDNA water sampling differed greatly in their frequency. (a) The number of taxa in different DNA sequence read categories. (b) The most common taxa defined by a naïve occupancy index defined as the proportion of sites where the species was detected. Notice that the three most detected mammals were domestic animals (cow, pig and dog). A full species list is provided in Supplementary Table S9.

### 3.3 Method comparison

Our review of conventional monitoring data from the Åmosen region yielded 263 wild vertebrate species. Of these, 29 species were deemed outliers as they were e.g., presumed locally extinct or were extremely rare visitors (supplementary table 11), resulting in a total of 234 final species for comparison with our CT and eDNA data (Figure 5a; Supplementary Table S11). We detected 115 wild species with eDNA and CT combined, including 20 fish, four amphibians, one snake, 23 mammals, and 67 birds (Figure 5, Table 1, Supplementary Table S9). Thus, the total number of wild species we detected during roughly 15 months of eDNA and four years of CT vertebrate monitoring comprise about half (115/234 = 49%) of the species observed in the region through decades of more conventional biomonitoring.

**FIGURE 5.**
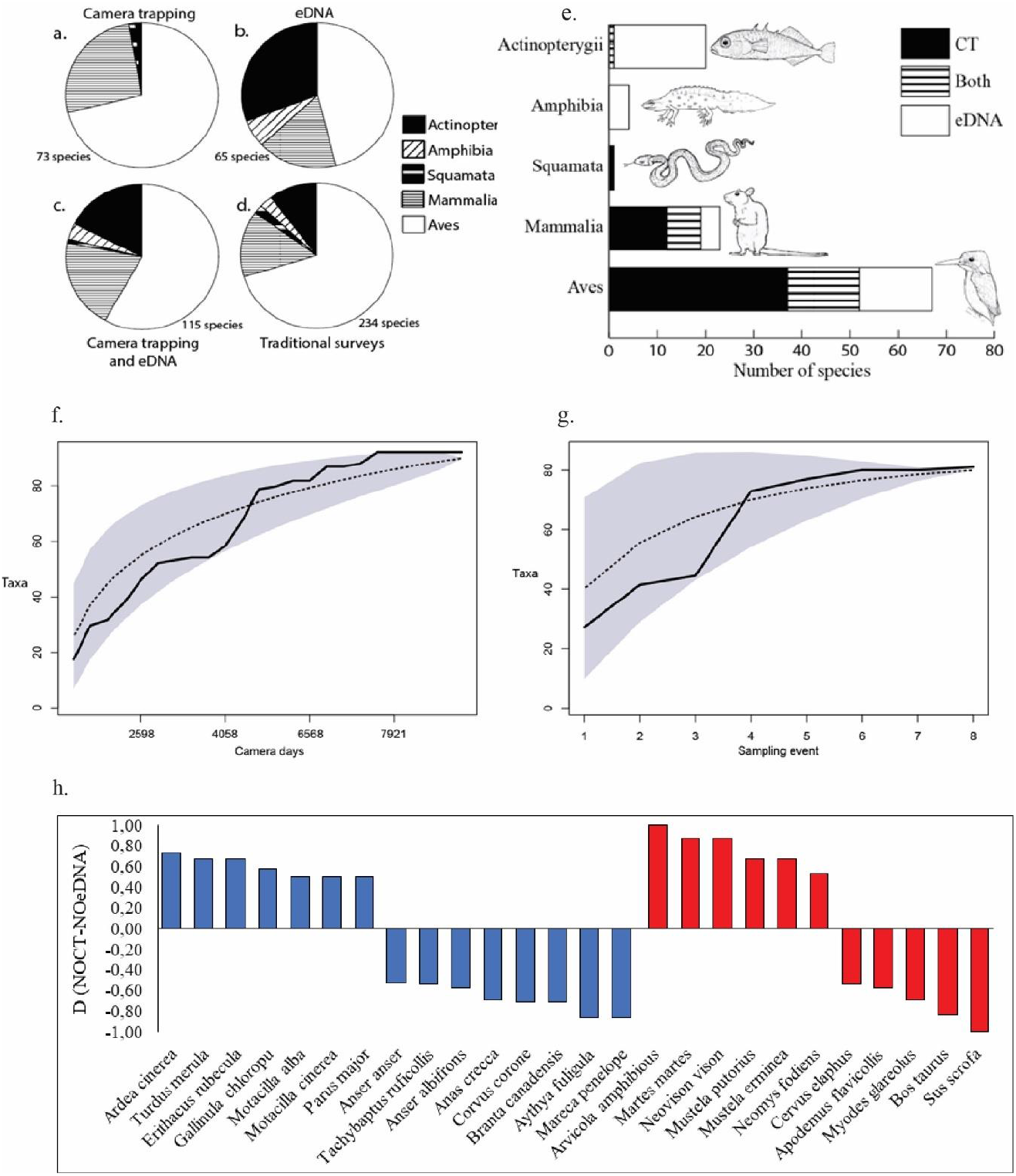
Evaluation of monitoring approach. (a) The number of taxa detected in Nature Park Åmosen for each vertebrate class as detected by camera trapping, (b) eDNA, (c) both methods, and (d) previous traditional surveys. (e) camera trapping and eDNA data compiled by each five verte-brate classes. (f-g) Species accumulation curves for vertebrate taxa detected in Åmosen by camera trapping and eDNA sampling. The black line are the detected taxa, stippled line is randomized accumulation curve estimated in specaccum (vegan package in R), and light grey shading is the 95% confidence intervals. h) Birds and mammals with large difference between camera trap naïve occupancy (NO_CT_) and eDNA naïve occupancy (NOeDNA). Species above or below the horizontal line are overrepresented in camera traps or eDNA, respectively. Illustrations by AMRH. A full species list is provided in Supplementary Table S9.

Only 30 species were detected with both eDNA and CTs, including one fish, seven mammals, and 15 birds (Figure 5 and 6). The aquatic eDNA detections were biased towards fish and amphibians, whereas CT detections were limited to mammals and birds, except for a single fish detection, which was a result of a grey heron (*Ardea cinerea*) catching a northern pike (*Esox lucius*) close to the camera (Figure 3b). The 115 species detected by CT and aquatic eDNA represented a large diversity in terms of body size, biomass, behaviour, life-history, habitat requirements and conservation status, including 19 species (16.5%) categorised as vulnerable, endangered, or critically endangered on the Danish Red List and seven species on the Natura 2000 list (EU Habitat Directive and/or Bird Directive) (Moeslund et al., 2019; Supplementary Figure 7; Supplementary Table S9).

**FIGURE 6.**
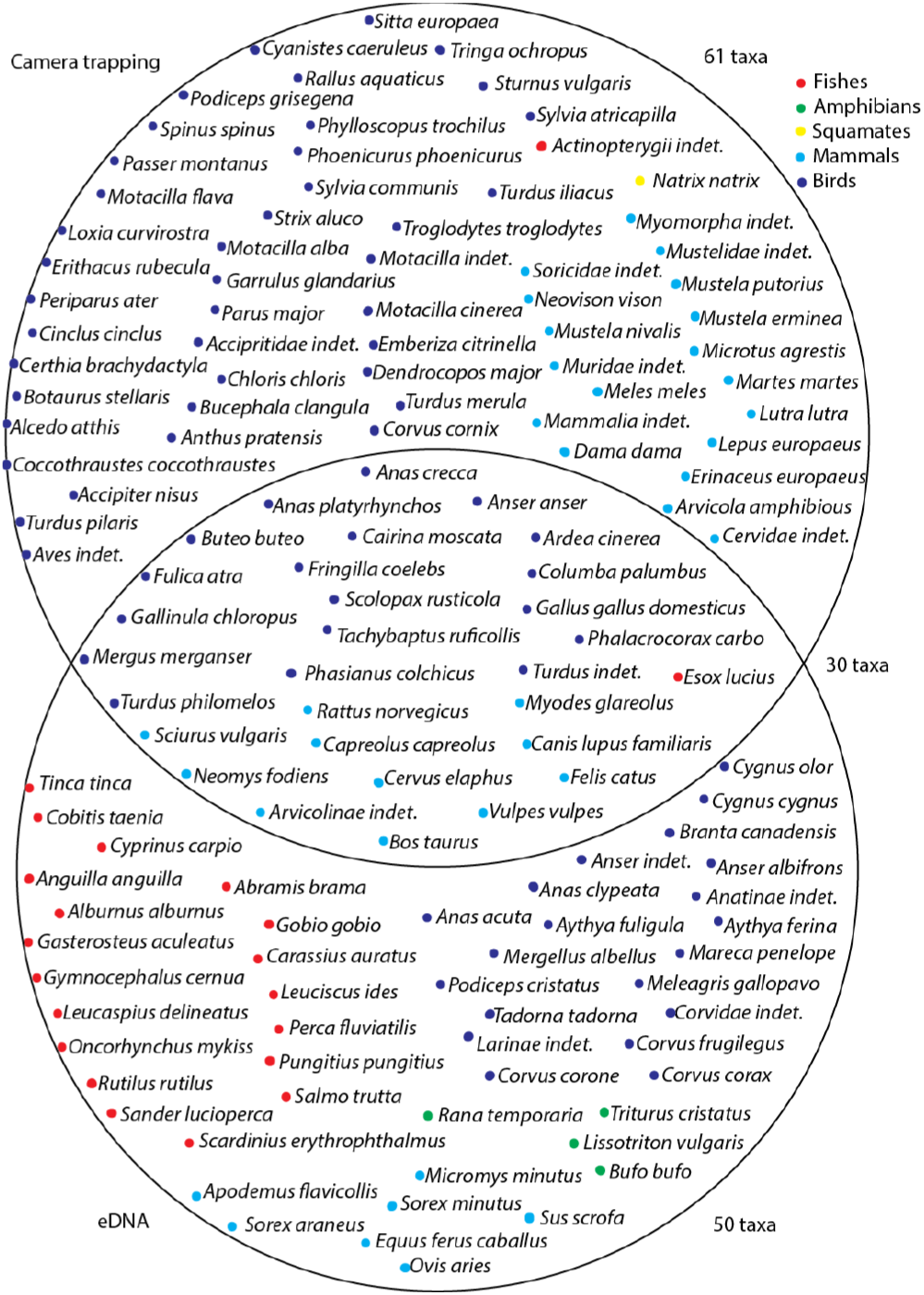
Overview of taxa found from camera trapping an eDNA. *Venn diagram* showing the overlap between the qualitative results obtained from camera trapping and eDNA metabarcoding of freshwater in Åmosen. Both methods detected 30 taxa, while 50 taxa only were detected by eDNA and 61 taxa only were detected by camera trapping.

## 4 Discussion

Our study demonstrates that CT and eDNA sampling can serve as complementary methods for a more holistic monitoring of the vertebrate fauna in temperate European wetland, nature park. We were able to verify the presence of 115 vertebrate species, which is nearly half of the total reported species from Åmosen (n=263) over the last 20 years (see suppl. Table S9 & S11). The taxa found with both CT and eDNA monitoring represent nearly 50 % of the eDNA taxa and around 46 % of the CT taxa, confirming the benefit of using the two methods in combination. These ratios are not much different from a terrestrial study of vertebrates in southwestern Australia combining soil eDNA and CT, with around half and one third of the total taxa occurring in eDNA and CTs, respectively (Ryan et al., 2021). It should, however, be considered that the CTs in the present study spanned across nearly four years and the eDNA monitoring only one year, making the comparison somewhat unbalanced.

With CTs, contrary to eDNA monitoring studies, the life stage of detected species can sometimes be determined, fx. juveniles of American mink (*Neovison vison*), mallard, pine marten, and stoat (*Mustela erminea*) detected by CTs in the present study. Foraging behavior was observed in several species including American mink, common wood pigeon, red fox, and white wagtail (*Motacilla alba*). On the other hand, some species can be hard to detect by CT due to their behaviour, life stage, or seasonal changes, potentially leading to biased results (Gotelli & Colwell, 2001). For such elusive species, parallel monitoring of eDNA is especially relevant for complementing CT and traditional monitoring methods. Amphibians can be hard to detect as they differ greatly in behaviour and appearance between life stages. Valentini et al., (2016) found that eDNA had a much higher detection rate of amphibians than traditional survey methods, provided that the sampling of eDNA is carried out while the amphibians are in their aquatic stage.

Monitoring of semi-aquatic animals, like Eurasian otter, can be problematic with CT (Lerone et al., 2015), and proved challenging when monitoring eDNA. Even though the primer set Mamm01 (Taberlet et al., 2018) was found to have no mismatches with otter DNA sequences obtained from NCBI GenBank database, we did not detect any eDNA from otter. Neither did we detect eDNA from any of the other seven species of mustelids (Figure 6, Supplementary table 9) detected by CTs even though comparison of the primers and the mtDNA target region in mustelid species did not show mismatches. Our CT data show that almost all mammal species were in contact with the freshwater stream at some point, and previous studies have shown that when terrestrial mammals drink from, or are otherwise in contact with, a water body, their DNA is often detectable in water samples (Matsubayashi et al., 2006; Rodgers and Mock, 2015; Ushio et al., 2017). Williams et al., (2017) found that even when only the snout of a pig was in contact with water, pig DNA could be detected in the water afterwards. Past studies have also shown difficulties in detecting eDNA from otter even when using a species-specific primer set (Thomsen et al., 2012; Andersen et al., 2018; Harper et al., 2019). Of all the seven Danish mustelid species (supplementary table 9), otters spend the most time in water (Baagøe and Jensen, 2007), but eDNA detection is likely challenging due to low populations sizes and/or nocturnal behaviour. Only 8% of eDNA monitoring studies far have targeted mammals (Tsuji et al., 2019), and future studies on monitoring eDNA from mammals could focus on how monitoring of semi-aquatic and fully terrestrial mammals can be optimized.

### Future perspectives

Efficient nature conservation and restoration increasingly requires non-invasive, cost-effective methods for monitoring biodiversity. The two methods used in the current study are already very useful for this task and they are still improving. CT is widely used for monitoring mammals, but while standardized protocols have been developed to estimate e.g., densities of large carnivores as well as factors affecting them (Havmøller et al., 2019), there is no single camera trap protocol that enables full insight into a vertebrate community, and camera trapping will unavoidably have taxon-specific biases (Burton et al., 2015). One of the most time-consuming factors with camera traps is data annotation, which is still largely done manually, although there are advances with annotation through machine learning (Whytock et al., 2021). In our study, 30% of all CT records contained an animal, while the rest were recordings triggered by moving water, vegetation, or heat spots from the sun. Camera traps are becoming cheaper and more efficient as the technology is developing. It is still considered a somewhat costly method, as equipment costs can be high, but the approach is comparatively cheap in the long-term. CT monitoring does not require experts in the field, but can instead rely on locals and volunteers, which has also been shown to broaden environmental awareness in local communities (Hönigsfeld-Adamic and Smole, 2011; Parsons et al., 2018).

Our study confirmed that the monitoring of eDNA is effective for monitoring the distribution and occurrence of both aquatic and semi-aquatic vertebrates as shown in other studies (Thomsen et al., 2012; Taberlet et al., 2018). Monitoring aquatic eDNA allowed for detection of all species of fish known from the area with the exception of a few rare species. Of the undetected species, grass carp (*Ctenopharyngodon idella*) and the Wels catfish (*Silurus glanis*) are known only from private ponds near the stream; the flounder (*Plectichtys flesus*) is mainly a marine species that occasionally migrates upstream to Tissø (Carl and Møller, 2012) and the burbut (*Lota lota*) went extinct in 1927 (Carl and Møller 2012). The only common species not detected was the Crucian carp (*Carsassius carassius*), a species mostly found in lentic waters, which might explain its absence in the river water samples. DNA metabarcoding is continuously being refined for more detailed multispecies detection (Creer et al., 2016), but we consider the aquatic eDNA metabarcoding method ready for large-scale monitoring of fish in European freshwater habitats. More terrestrial mammals might have been detected if eDNA from soil, dung or air samples had been included as well (Leempoel et al., 2020; Sales et al., 2020; Lynggaard et al., 2021; van der Heyde et al., 2021).

Like many other Danish nature parks and national parks, Nature Park Åmosen is a mosaic of cultural landscapes and more natural habitats mixed with human installations, roads, cities and agriculture. As demonstrated in the present study shows these parks can host a variety of wildlife, especially in small pockets of old forest and around near-natural rivers. Such a biodiversity hot-spot is our sampling site Kattrup, with almost twice as many species as the other sites. This is also where we first found the otter, which is extremely rare on the island of Zealand. Combining CTs and eDNA metabarcoding could be an efficient future means for vertebrate biodiversity monitoring in wetlands and other wildlife habitats.

## Acknowledgements

Special thanks go to all the people who helped in making this project possible. To 15. Juni Fonden for financial support, to Nature Park Åmosen for allowing research in the area; to Hans Henrik Erhardi and The Danish Nature Agency, Vestsjælland for logistics and introduction to the area; to Hans Erik Svart for providing to first camera traps and for introduction to the Zealand otter. Thanks to the private landowners at Kattrup Gods especially Jens Hansen and Jakob Bak, for their help and cooperation. Special thanks to Jan Bolding Kristensen, Mikkel H. Post and Hans J. Baagøe who helped verify bird and mammal identifications and to Henrik Carl for help with fish data.

## Notes

### Competing Interest Statement

The authors have declared no competing interest.

