## Supplemental material for "Holistic monitoring of freshwater and terrestrial vertebrates by camera trapping and environmental DNA"

### Supporting text with details on methods applied

#### Supporting text A:

For the qPCR (quantitative polymerase chain reaction) setup, we used tagged versions of the MiFish primer set (MiFish-U-F: 5'-GTCGGTAAAACTCGTGCCAGC-3' and MiFish-U-R: 5'-CATAGTGGGGTATCTAATCCCAGTTTG-3') that targets a 163-185 bp part of the mitochondrial DNA 12S rRNA gene (Miya al. 2015). Many of the extracted samples looked brownish and dirty.

Discoloration can indicate the presence of humic substances in the extraction, which can inhibit the subsequent PCR procedure (Wilson 1997). Dilution of the extracted DNA material can minimize this inhibition (Jane 2015, Wilcox et al. 2015, McKee 2015, Spear et al. 2015). To determine inhibitors are present in the extractions, we made a dilution series (1:1, 1:8 and 1:32) of three different samples with different degrees of discoloration and performed a qPCR analysis on an Agilent Stratagene Mx3005P machine. This made it possible to check if diluted extraction more easily could amplify the environmental DNA target fragment, and thereby help avoid inhibition in the later PCR metabarcoding setup. Extracted samples were diluted in $ddH_2O$. Each qPCR tube was prepared in 25 µL total volume reactions comprising: 1 µL MiFish primer (forward and reverse) (10 mM each), 0.2 µL TaqGold polymerase (5 U/µL), 2 µL dNTPs (2 mM/ dNTP), 2 µL $MgCl_2$ (25 mM), 2.5 µL AmpliTaq buffer (x10), 0.25 µL BSA (20 mg/mL), 13.05 µL $ddH_2O$, 1 µL SYBR green mix [prepared from 1 part SYBR Green (Invitrogen cat. no. S7563), 4 parts ROX reference dye (Invitrogen cat. no. 12223-012), and 2000 parts DMSO (Invitrogen)], and 3 µL diluted DNA-template – i.e. dilution levels being 1:1, 1:8 and 1:32 of original extracted water sample. The thermocycling parameters were set at 95 °C for 10 minutes, followed by 45 cycles of (94 °C in 30 s, 50 °C for 30 s and 72 °C for 1 minute), with collecting of flourescence at end-point in extension phase. The qPCR with diluted extraction indicated the highest dilution at 1:32 performed best, by amplifying at the earliest cycle of quantification. Based on this, we diluted all the extracted samples into 1:32 for the PCR metabarcoding setup to prevent or minimize inhibition.

**Supporting text B:**

The PCR for metabarcoding was performed in three replicates per sample. Using the primer set Mamm01 (mamm01_F:5’-CCGCCCGTCACCCTCCT-3’, mamm01_R: 5’-GTAYRCTTACCWTGTTACGAC-3’), which is optimized for mammals (Taberlet et al., 2018), and the primer set MiFish-U (MiFish-U_F: 5’-GTCGGTAAAACTCGTGCCAGC-3’, MiFish-U_R: 5’-

CATAGTGGGGTATCTAATCCCAGTTTG-3'), which is optimized for bony fishes (Miya et al., 2015). Each PCR was done in a 25 µL volume, comprised 13.05 µL $ddH_2O$, 2.0 µL dNTPs (2.0 mM/dNTP), 2.5 µL GeneAmp® 10x PCR Gold Buffer, 1.5 µL MgCl2 (25 mM), 0.25 BSA (20 mg/µL), 0.2 µL AmpliTaq GoldTM DNA Polymerase (5 U/µL), 1.25 µL of each primer (10 mM) (forward and reverse), and 3 µL extracted DNA (diluted 1:32). A 1:32 dilution of the extracted DNA was preferred, based on a qPCR analysis testing for inhibition at three different dilution levels (1:1, 1:8, and 1:32). The qPCR showed that the highest dilution level (1:32) yielded the earliest onset of amplification (see supplementary material text A). Thermocycling parameters of the PCR with the MiFish-U primer set were set to an initial 5-minute denaturation at 95ºC, followed by 40 cycles of denaturation at 94ºC for 30 s, annealing at 55ºC for 30 s and extension at 72ºC for 90 s, and then a final extension at 72ºC for 10 minutes. For the Mamm01 primer set, PCR thermocycling parameters were set to an initial 5-minute denaturation at 95ºC, followed by 37 cycles of denaturation at 94ºC for 30 s, annealing at 53ºC for 30 s and extension at 72ºC for 90 s, and then a final extension for 72ºC for 10 minutes, as in previous protocols (Sigsgaard et al., 2017). Setup of PCR for metabarcoding is outlined in supplementary table S1-S8.

**Supporting text C:**

From each of the 12 libraries (2*3 replicates for MiFish-U and 2*3 replicates for Mamm01) (supplementary material table S1-S8) we pooled 10 µL to a total of 120 µL. The 120 µL was then purified using the MinElute® PCR purification kit (cat. nos. 28006), following the supplied protocol with an additional centrifuge step for 60 s at 10,000 G after discarding flow-through, and a heat step at 37ºC for 10 minutes, before the spin column was added to a new 1.5 mL LoBind Eppendorf tube. We added 20 µL EB buffer, heated to 37ºC for 10 minutes and centrifuged for 1 minute at 10,000 G, and repeated the addition of 20 µL EB buffer and the heat step, giving a total of 40 µL per pooled sample.

**Table. S1.** PCR 37 with the Mamm01 primer setup. The yellow plate column number (plt.c No) and plate row number (plt.r No) indicates the individual tag (Tag no) for each sample, the orange color denotes the different pooled samples, except for NegC (negative control) and Mock (positive control), and the blue the three replicates (roman numerals) for each sample. Each sample have had their own tag number, which was the same number (e.g. Tag01) for both forward and reverse primer. 'Lib.no' indicates lettercode for pooled libraries A to C. See table S7 for list of tags used for primers. Sample abbreviations (Sampl Abbr) are explained in table S8.

| | plt.c No | 1 | 2 | 3 | | 4 | 5 | 6 | | 7 | 8 | 9 |
|---|---|---|---|---|---|---|---|---|---|---|---|---|
| plt.r No | Sampl Abbr | Tag no | Tag no | Tag no | Sampl Abbr | Tag no | Tag no | Tag no | Sampl Abbr | Tag no | Tag no | Tag no |
| A | Und2 | T01 | T01 | T01 | Skel9 | T09 | T09 | T09 | Kat12 | T17 | T17 | T17 |
| B | Und3 | T02 | T02 | T02 | Skel12 | T10 | T10 | T10 | Str9-17 | T18 | T18 | T18 |
| C | Und5 | T03 | T03 | T03 | Kat9-17 | T11 | T11 | T11 | Str10-17 | T19 | T19 | T19 |
| D | Und9 | T04 | T04 | T04 | Kat10-17 | T12 | T12 | T12 | Str1 | T20 | T20 | T20 |
| E | Und12 | T05 | T05 | T05 | Kat2 | T13 | T13 | T13 | Str2 | T21 | T21 | T21 |
| F | Skel2 | T06 | T06 | T06 | Kat3 | T14 | T14 | T14 | Str3 | T22 | T22 | T22 |
| G | Skel3 | T07 | T07 | T07 | Kat5 | T15 | T15 | T15 | Mock | T30 | T30 | T30 |
| H | Skel5 | T08 | T08 | T08 | Kat9 | T16 | T16 | T16 | NegC | T31 | T31 | T31 |
| | Repl No | I | II | III | | I | II | III | | I | II | III |
| | Lib No | A | B | C | | A | B | C | | A | B | C |

**Table. S2.** PCR 38 with the Mamm01 primer setup. The yellow plate column number (plt.c No) and plate row number (plt.r No) indicates the individual tag (Tag no) for each sample, the orange color denotes the different pooled samples, except for NegC (negative control) and Mock (positive control), and the blue the three replicates (roman numerals) for each sample. Each sample have had their own tag number, which was the same number (e.g. Tag01) for both forward and reverse primer. 'Lib.no' indicates lettercode for pooled libraries E to F. See table S7 for list of tags used for primers. Sample abbreviations (Sampl Abbr) are explained in table S8.

| | plt.c No | 1 | 2 | 3 | | 4 | 5 | 6 | | 7 | 8 | 9 |
|---|---|---|---|---|---|---|---|---|---|---|---|---|
| plt.r No | Sampl Abbr | Tag no | Tag no | Tag no | Sampl Abbr | Tag no | Tag no | Tag no | Sampl Abbr | Tag no | Tag no | Tag no |
| A | Str2 | T01 | T01 | T01 | Tisin12 | T09 | T09 | T09 | Bøs5 | T17 | T17 | T17 |
| B | Str9 | T02 | T02 | T02 | Tisout2 | T10 | T10 | T10 | Bøs9 | T18 | T18 | T18 |
| C | Str12 | T03 | T03 | T03 | Tisout3 | T11 | T11 | T11 | Bøs12 | T19 | T19 | T19 |
| D | Mad | T04 | T04 | T04 | Tisout5 | T12 | T12 | T12 | NegC | T30 | T30 | T30 |
| E | Tisin2 | T05 | T05 | T05 | Tisout9 | T13 | T13 | T13 | Mock | T31 | T31 | T31 |
| F | Tisin3 | T06 | T06 | T06 | Tisout12 | T14 | T14 | T14 | | | | |
| G | Tisin5 | T07 | T07 | T07 | Bøs2 | T15 | T15 | T15 | | | | |
| H | Tisin9 | T08 | T08 | T08 | Bøs3 | T16 | T16 | T16 | | | | |
| | Repl No | I | II | III | | I | II | III | | I | II | III |
| | Lib No | E | D | F | | E | D | F | | E | D | F |

**Table. S3.** PCR 21 with the MiFish-U primer setup. The yellow plate column number (plt.c No) and plate row number (plt.r No) indicates the individual tag (Tag no) for each sample, the orange color denotes the different pooled samples, except for NegC (negative control) and Mock (positive control), and the blue the three replicates (roman numerals) for each sample. Each sample have had their own tag number, which was the same number (e.g. Tag01) for both forward and reverse primer. 'Lib.no' indicates lettercode for pooled libraries G to I. See table S7 for list of tags used for primers. Sample abbreviations (Sampl Abbr) are explained in table S8.

| | plt.c No | 1 | 2 | 3 | | 4 | 5 | 6 |
|---|---|---|---|---|---|---|---|---|
| plt.r No | Sampl Abbr | Tag no | Tag no | Tag no | Sampl Abbr | Tag no | Tag no | Tag no |
| A | Und9 | T45 | T45 | T45 | Str9 | T53 | T53 | T53 |
| B | Und12 | T46 | T46 | T46 | Str12 | T54 | T54 | T54 |
| C | Skel9 | T47 | T47 | T47 | NegC | T55 | T55 | T55 |
| D | Skel12 | T48 | T48 | T48 | Mock | T56 | T56 | T56 |
| E | Kat9-17 | T49 | T49 | T49 | | | | |
| F | Kat9 | T50 | T50 | T50 | | | | |
| G | Kat12 | T51 | T51 | T51 | | | | |
| H | Str9-17 | T52 | T52 | T52 | | | | |
| | Repl No | I | II | III | | I | II | III |
| | Lib No | G | H | I | | G | H | I |

**Table. S4.** PCR 22 with the MiFish-U primer setup. The yellow plate column number (plt.c No) and plate row number (plt.r No) indicates the individual tag (Tag no) for each sample, the orange color denotes the different pooled samples, except for NegC (negative control) and Mock (positive control), and the blue the three replicates (roman numerals) for each sample. Each sample have had their own tag number, which was the same number (e.g. Tag01) for both forward and reverse primer. 'Lib.no' indicates lettercode for pooled libraries G to I. See table S7 for list of tags used for primers. Sample abbreviations (Sampl Abbr) are explained in table S8.

| | plt.c No | 1 | 2 | 3 | | 4 | 5 | 6 |
|---|---|---|---|---|---|---|---|---|
| plt.r No | Sampl Abbr | Tag no | Tag no | Tag no | Sampl Abbr | Tag no | Tag no | Tag no |
| A | Mad9 | T45 | T45 | T45 | Mock | T53 | T53 | T53 |
| B | Tisin9 | T46 | T46 | T46 | | | | |
| C | Tisin12 | T47 | T47 | T47 | | | | |
| D | Tisout9 | T48 | T48 | T48 | | | | |
| E | Tisout12 | T49 | T49 | T49 | | | | |
| F | Bøs9 | T50 | T50 | T50 | | | | |
| G | Bøs12 | T51 | T51 | T51 | | | | |
| H | NegC | T52 | T52 | T52 | | | | |
| | Repl No | I | II | III | | I | II | III |
| | Lib No | J | K | L | | J | K | L |

**Table S5.** Measured concentrations on Qubit for all libraries (Lib No) after pooling of samples and MinElute purification in order to get a sufficiently high concentration (ng) to build libraries.

| Lib No | Qubit concentration (ng/µL) | Amount of sample (µL) added in order to get 250 ng | EB Buffer added to get final volume of 60 µL |
|---|---|---|---|
| A | 6.98 | 35.8 | 24.2 |
| B | 6.68 | 37.4 | 22.6 |
| C | 7.60 | 32.9 | 27.1 |
| D | 9.44 | 26.5 | 33.5 |
| E | 8.54 | 29.3 | 30.7 |
| F | 9.56 | 26.2 | 33.8 |
| G | 33.60 | 14.9 | 45.1 |
| H | 40.80 | 12.3 | 47.7 |
| I | 32.80 | 15.2 | 44.8 |
| J | 24.00 | 20.8 | 39.2 |
| K | 23.80 | 21.0 | 39.0 |
| L | 24.80 | 20.2 | 39.8 |

**Table S6.** Information needed to pool samples in order to send to the High-throughput DNA Sequencing Centre at the University of Copenhagen. The Tapestation results were used for evaluating the fragment length. Abbreviations used above columns are: 'Lib No' library number; 'TiNm' TruSeq adapter-index name; 'Aseq' Adapter sequence; 'CTS' Conc (ng/µL) TS, Tapestation; 'CQb' Conc (ng/µL), Qubit; 'Psz' Peak size (bp) measured by Tapestation; 'CPk' Conc.of peak (ng/µL), measured by Tapestation; 'VPur' Vol. of purified library (µL) after Qubit and Tapestation; 'Insz' Insert size (bp), incl. primers and tags and adaptor indexed ligated; 'LCo' Lowest conc. (nM); 'c' (Concentration in ng/ul) / (660 g/mol * average library size in bp) * 10^6); 'Ac' Average c (nM); 'V1' A volume between 6 µL and volume of purified library; 'V2' Volume (µL) to be used for pooling; 'Psz Ntg' Product size (bp) without tags.

| Lib No | TiNm | Aseq | CTS | CQb | Psz | CPk | VPur | Insz | LCo | c | Ac | V1 | V2 | Psz Ntg |
|---|---|---|---|---|---|---|---|---|---|---|---|---|---|---|
| A | AR001 | ATCACG(A) | 56.5 | 4.4 | 530 | 20.5 | 27 | 237 | 27.5 | 28.4 | 29 | 9 | 9 | 97 |
| B | AR003 | TTAGGC(A) | 228.0 | 4.5 | 537 | 76.5 | 27 | 237 | 27.5 | 28.5 | 29 | 9 | 9 | 97 |
| C | AR008 | ACTTGA(A) | 66.9 | 5.2 | 537 | 13.2 | 27 | 237 | 27.5 | 33.0 | 29 | 9 | 8 | 97 |
| D | AR009 | GATCAG(A) | 76.5 | 4.3 | 512 | 25.4 | 27 | 237 | 27.5 | 27.7 | 29 | 9 | 9 | 97 |
| E | AR010 | TAGCTT(A) | 125.0 | 4.3 | 525 | 95.7 | 27 | 237 | 27.5 | 27.5 | 29 | 9 | 9 | 97 |
| F | AR011 | GGCTAC(A) | 58.2 | 4.4 | 497 | 43.1 | 27 | 237 | 27.5 | 28.1 | 29 | 9 | 9 | 97 |
| G | AR002 | CGATGT(A) | 12.1 | 5.0 | 622 | 6.9 | 27 | 358 | 17.9 | 21.3 | 23 | 9 | 8 | 218 |
| H | AR004 | TGACCA(A) | 11.1 | 5.0 | 622 | 6.9 | 27 | 358 | 17.9 | 21.2 | 23 | 9 | 8 | 218 |
| I | AR005 | ACAGTG(A) | 8.9 | 4.2 | 591 | 5.4 | 27 | 358 | 17.9 | 17.9 | 23 | 9 | 9 | 218 |
| J | AR006 | GCCAAT(A) | 16.4 | 6.6 | 582 | 9.5 | 27 | 358 | 17.9 | 28.1 | 23 | 9 | 6 | 218 |
| K | AR007 | CAGATC(A) | 13.3 | 5.9 | 607 | 8 | 27 | 358 | 17.9 | 25.1 | 23 | 9 | 6 | 218 |
| L | AR012 | CTTGTA(A) | 11.7 | 5.6 | 617 | 7 | 27 | 358 | 17.9 | 23.6 | 23 | 9 | 7 | 218 |

**Table S7.** List of tags and their corresponding nucleotide sequence used for each primer set.

| Tag number | Tag nucleotide sequence | Primerset |
|---|---|---|
| T01 | AACAAC | Mamm01 |
| T02 | AACCGA | Mamm01 |
| T03 | CCGGAA | Mamm01 |
| T04 | AGTGTT | Mamm01 |
| T05 | CCGCTG | Mamm01 |
| T06 | AACGCG | Mamm01 |
| T07 | GGCTAC | Mamm01 |
| T08 | TTCTCG | Mamm01 |
| T09 | TCACTC | Mamm01 |
| T10 | GAACTA | Mamm01 |
| T11 | CCGTCC | Mamm01 |
| T12 | AAGACA | Mamm01 |
| T13 | CGTGCG | Mamm01 |
| T14 | GGTAAG | Mamm01 |
| T15 | ATAATT | Mamm01 |
| T16 | CGTCAC | Mamm01 |
| T17 | TTGAGT | Mamm01 |
| T18 | AAGCAG | Mamm01 |
| T19 | TTGCAA | Mamm01 |
| T20 | CACGTA | Mamm01 |
| T21 | TAACAT | Mamm01 |
| T22 | TGCGTG | Mamm01 |
| T30 | GTACAC | Mamm01 |
| T31 | AAGTGT | Mamm01 |
| T45 | CTATAA | MiFi-U |
| T46 | AATGAA | MiFi-U |
| T47 | CGAATC | MiFi-U |
| T48 | AGAGAC | MiFi-U |
| T49 | TTCGGA | MiFi-U |
| T50 | CGACGT | MiFi-U |
| T51 | CTCATG | MiFi-U |
| T52 | TGTATA | MiFi-U |
| T53 | ACAACC | MiFi-U |
| T54 | TCAGAG | MiFi-U |
| T55 | GTAGTG | MiFi-U |
| T56 | AGCACT | MiFi-U |

**Table S8.** Overview of samples. Study sites, sampling date, number of replicates and volume of filtered water. Laboratory work by MM was with the MiFish-U primerset (Miya et al, 2015). Sampling location abbreviations and filtered volume of water sample. Labwork was performed by Anne Marie Rubæk Holm (AMRH) and Malene Månson (MM). Sampling locations are abbreviated as: 'Bøs' Bøstrup; 'Kat' Kattrup; 'Mad' Madesø; 'Skel' Skellingsted; 'Str' Stridsmølle; 'Tisin' Tissø inlet; 'Tisout' Tissø outlet; 'Und' Undløse.

| Sampl | Date | No. | Filtered | Labwork |
|---|---|---|---|---|

| Abbreviation | | replicates | volume (mL) |
|---|---|---|---|
| Bøs | 2018.Dec.13 | 3 | 3 x 400 AMRH |
| Bøs | 2018.Feb.27 | 3 | 3 x 350 MM |
| Bøs | 2018.Mar.28 | 3 | 3 x 500 MM |
| Bøs | 2018.May.09 | 3 | 3 x 300 MM |
| Bøs | 2018.Sep.04 | 3 | 3 x 60 AMRH |
| Kat | 2017.Oct.26 | 2 | 2 x 300 MM |
| Kat | 2017.Sep.27 | 2 | 2 x 500 AMRH |
| Kat | 2018.Feb.27 | 3 | 3 x 200 MM |
| Kat | 2018.Mar.28 | 3 | 3 x 350 MM |
| Kat | 2018.May.09 | 3 | 3 x 600 MM |
| Kat | 2018.Sep.04 | 3 | 3 x 1000 AMRH |
| Mad | 2017.Sep.27 | 2 | 2 x 500 AMRH |
| Skel | 2018.Dec.13 | 3 | 3 x 400 AMRH |
| Skel | 2018.Feb.27 | 3 | 3 x 200 MM |
| Skel | 2018.Mar.28 | 3 | 3 x 400 MM |
| Skel | 2018.May.09 | 3 | 420/480/480 MM |
| Skel | 2018.Sep.04 | 3 | 3 x 600 AMRH |
| Str | 2017.Oct.26 | 2 | 2 x 300 MM |
| Str | 2017.Sep.17 | 2 | 2 x 500 AMRH |
| Str | 2018.Dec.13 | 3 | 3 x 400 AMRH |
| Str | 2018.Dec.13 | 3 | 3 x 400 AMRH |
| Str | 2018.Feb.27 | 3 | 240/230/205 MM |
| Str | 2018.Jan.31 | 3 | 3 x 300 MM |
| Str | 2018.Mar.28 | 3 | 3 x 350 MM |
| Str | 2018.May.09 | 3 | 480/600/540 MM |
| Str | 2018.Sep.04 | 3 | 3 x 600 AMRH |
| Tisin | 2018.Dec.13 | 3 | 3 x 400 AMRH |
| Tisin | 2018.Feb.27 | 2 | 3 x 200 MM |
| Tisin | 2018.Mar.28 | 3 | 500/450/450 MM |
| Tisin | 2018.May.09 | 3 | 3 x 500 MM |
| Tisin | 2018.Sep.04 | 3 | 3 x 600 AMRH |
| Tisout | 2018.Dec.13 | 3 | 3 x 300 AMRH |
| Tisout | 2018.Feb.27 | 2 | 3 x 400 MM |
| Tisout | 2018.Mar.28 | 3 | 3 x 400 MM |
| Tisout | 2018.May.09 | 3 | 3 x 500 MM |
| Tisout | 2018.Sep.04 | 3 | 3 x 300 AMRH |
| Und | 2018.Dec.13 | 3 | 3 x 600 AMRH |
| Und | 2018.Feb.27 | 3 | 3 x 270 MM |
| Und | 2018.Mar.28 | 3 | 500/500/450 MM |
| Und | 2018.May.09 | 3 | 780/720/720 MM |
| Und | 2018.Sep.04 | 3 | 3 x 1000 AMRH |

**Supporting text D:**

**Results of mainly local interest.**

Seven fish families were found in total, and at five of the seven sites, all these families were represented. At the two remaining sites, Bøstrup and Tissø inlet, Salmonidae and Cobitidae were absent, respectively. At each of the study sites, the number of fish species detected by eDNA varied between 12 and 19 species, with Kattrup having the largest diversity and Undløse the lowest (Table 1, Suppl. Table S9).

The only detected reptile, grass snake (*Natrix natrix*), was found at Skellingsted. Amphibians were detected at six of the eight study sites. The northern crested newt was detected at Stridsmølle and Undløse, where the smooth newt was detected at Bøstrup and Tissø inlet (Suppl. Table 1).

Mammals were found present at all eight sites via both CT and eDNA and varied in richness from four to18 species per site. Hedgehog (*Erinaceus europaeus*) and European hare (*Lepus europaeus*) were only detected in Kattrup. None of the three Soricomorpha were detected south of Tissø (neither at Bøstrup or Tissø outlet). Four and six different mammals were detected by eDNA alone at Tissø inlet and Tissø outlet, respectively, as no CT were deployed at these two sites. Six mammal species were detected at Bøstrup with both CT and eDNA (Table 1, Suppl. Table S9). Birds were found present at all eight locations with both CT and eDNA. The number of detected species per site varied from 18 to 47. Kattrup showed the highest diversity of bird species (47 species), while Stridsmølle and Undløse yielded roughly half this richness (23 species at each site). Sites south of Tissø (Bøstrup and Tissø outlet) held the lowest number of bird orders (4 and 5 species, respectively), with Charadriiformes, Columbiformes, Accipitriformes, Coraciiformes, Galliformes, Piciformes, and Strigiformes being absent (Suppl. Table S9, Suppl. Figure 6). At none of the sites were all the 12 orders of birds known from the area represented. Bøstrup, Tissø inlet, and Tissø outlet had 11 to12 different waterfowl (Anseriformes) at each site. Kattrup had the largest number of passerines (Passeriformes, 25 species).

Northern crested newt and smooth newt were not detected at the same sites (Supp. Table S9). Smooth newt was only detected at Bøstrup and Tissø inlet, while northern crested newt was detected at Stridsmølle and Undløse. The smooth newt often inhabits meadows, forests, and uncultivated areas, whereas the northern crested newt to a larger degree is dependent on forests and areas with settlements (Fog *et al.*, 2001a, 2001b). Their preferred habitats were confirmed by the eDNA monitored here (Table 1, Suppl. Table S9), but additional research is needed to verify if this is a general trend. Monitoring of these amphibians may turn out to be difficult as the eDNA of stream dwelling salamanders have been shown to be detectable only within few meters of the source animal (Pilliod *et al.* 2014).

The number of mammals at each study site varied a lot, from four (Tissø outlet) to 18 species (Kattrup) (Table 1). The 23 wild mammal species found in total accounts for 68 % of the mammals known from this area from traditional surveys. As the mammals recorded in Åmosen inhabit a wide variety of strata (e.g., fossorial, terrestrial, aquatic, volant or arboreal) the effort needed to efficiently monitor their diversity and distribution need further diversification, as some species was almost never be detected by the CT or eDNA sampling design in our study. The European mole (*Talpa europaea*) are common but fossorial, and six different bats (Vespertilionidae) forage in the evenings outside the hours where eDNA sampling was done, so the absence of these animals from the recorded vertebrates might reflect their ecology and strata they inhabit (Baagøe and Jensen, 2007). Excluding species, that alludes to our monitoring design, raises our diversity record percentage from 68% to 85 %.

The overall detection of birds varied from 18 (Bøstrup) to 47 (Kattrup). The number of birds found was likely caused by the differences in habitat composition i.e. Kattrup having many, tall, old trees. Tissø inlet and outlet had the highest number of waterfowls, and this match these two areas being an oasis for waterfowl species. Several Passerine birds were detected at Kattrup (n = 25) compared to the remaining sites with 2-11 records. Kattrup has forest on one side of the stream with cover, and a mix of habitat types with more open areas on the other side and makes this habitat diverse for species that require both hiding and foraging places in their habitat (European Commission, 2007; Naturstyrelsen, 2013, 2014).

**Table S9.** Overview of results from camera trapping and eDNA in Åmosen. For each taxon, the number of camera events (CE) (sightings of animal(s) of same species with at least a 30-minute interval), the number of CE of a given taxon per 100 camera days (relative abundance index; RAI) and naïve occupancy (number of sites/samples that are positive to taxon presence divided by the total number of sites/samples) of CT data (six CT sampling sites, and 18 sampling locations in total), together with number of reads and Naïve occupancy from eDNA data (seven sampling sites, and 40 samples in total) is given. DOM = domestic species. Danish Red List status: NT = Near threatened, VU = Vulnerable, EN = Endangered, and CR = Critically endangered (Moeslund et al., 2019). Natura 2000 protection: (N2000) (Naturstyrelsen, 2012). Note that eDNA was not sampled at Bromølle, and CTs were not placed at Tissø inlet and Tissø outlet. Abbreviations in table represents: Camera trapping (CE), environmental DNA (eDNA), relative abundance index (RAI), number of sites (ns), number of locations (nl), number of reads (nr), number of samples (nspl). Sample sites are abbreviated: Bromølle (Bro), Bøstrup (Bøs), Kattrup (Kat), Skellingsted (Ske), Stridsmølle (Str), Tissøinlet (Tisin), Tissøoutlet (Tisou), Undløse (Und).

| **Class/Family** | | **Camera trapping** | | | | **Environmental DNA** | | | | | | | | | | |
|---|---|---|---|---|---|---|---|---|---|---|---|---|---|---|---|---|
| **Taxon** | **Status** | **CE** | **RAI** | **ns** | **nl** | **Reads** | **ns** | **nspl** | **Bro** | **Bøs** | **Kat** | **Ske** | **Str** | **Tisin** | **Tisou** | **Und** |
| **Actinopterygii** | | | | | | | | | | | | | | | | |
| Anguillidae | | | | | | | | | | | | | | | | |
| *Anguilla anguilla* | CR | | | | | 160074 | 1.0 | 0.6 | | x | x | x | x | x | x | x |
| Cobitidae | | | | | | | | | | | | | | | | |
| *Cobitis taenia* | N2000 | | | | | 3667 | 0.9 | 0.2 | | x | x | x | x | | x | x |
| Cyprinidae | | | | | | | | | | | | | | | | |
| *Abramis brama* | | | | | | 170362 | 0.9 | 0.3 | | x | x | x | x | x | x | |
| *Alburnus alburnus* | | | | | | 95164 | 0.7 | 0.2 | | x | x | | x | x | x | |
| *Carassius auratus* | | | | | | 1523 | 0.3 | 0.1 | | | | x | x | | | |
| *Cyprinus carpio* | | | | | | 622 | 0.7 | 0.2 | | | x | x | x | | x | x |
| *Gobio gobio* | | | | | | 148211 | 0.7 | 0.6 | | | x | x | x | x | | x |
| *Leucaspius delineatus* | | | | | | 198648 | 0.9 | 0.4 | | x | x | x | x | x | | x |
| *Leuciscus idus* | | | | | | 556969 | 1.0 | 0.5 | | x | x | x | x | x | x | x |
| *Rutilus rutilus* | | | | | | 2541591 | 1.0 | 1.0 | | x | x | x | x | x | x | x |
| *Scardinius erythrophthalmus* | | | | | | 16458 | 1.0 | 0.2 | | x | x | x | x | x | x | x |
| *Tinca tinca* | | | | | | 193480 | 0.9 | 0.4 | | x | x | x | x | x | x | |
| Esocidae | | | | | | | | | | | | | | | | |
| *Esox lucius* | | 1 | 0.0 | 0.2 | 0.1 | 305488 | 1.0 | 0.8 | | x | x | x | x | x | x | x |

| | | | | | | | | | | | | | | | | |
|---|---|---|---|---|---|---|---|---|---|---|---|---|---|---|---|---|
| Gasterosteidae | | | | | | | | | | | | | | | | |
| *Gasterosteus aculeatus* | | | | | | 205202 | 0.7 | 0.3 | | x | x | x | | x | x | |
| *Pungitius pungitius* | | | | | | 346094 | 1.0 | 0.8 | | x | x | x | x | x | x | x |
| Percidae | | | | | | | | | | | | | | | | |
| *Gymnocephalus cernua* | | | | | | 753260 | 0.7 | 0.4 | | x | x | x | | x | x | |
| *Perca fluviatilis* | | | | | | 1776745 | 1.0 | 0.9 | | x | x | x | x | x | x | x |
| *Sander lucioperca* | | | | | | 12886 | 0.4 | 0.1 | | | x | | | x | x | |
| Salmonidae | | | | | | | | | | | | | | | | |
| *Oncorhynchus mykiss* | | | | | | 1866 | 0.6 | 0.2 | | | x | x | x | | x | |
| *Salmo trutta* | | | | | | 59469 | 0.9 | 0.5 | | | x | x | x | x | x | x |
| **Amphibia** | | | | | | | | | | | | | | | | |
| Bufonidae | | | | | | | | | | | | | | | | |
| *Bufo bufo* | | | | | | 2679 | 0.7 | 0.2 | | x | x | x | x | | | x |
| Ranidae | | | | | | | | | | | | | | | | |
| *Rana temporaria* | NT | | | | | 344 | 0.4 | 0.1 | | | | | x | | x | x |
| Salamandridae | | | | | | | | | | | | | | | | |
| *Lissotriton vulgaris* | | | | | | 216 | 0.3 | 0.1 | | x | | | | x | | |
| *Triturus cristatus* | N2000 | | | | | 4725 | 0.3 | 0.1 | | | | | x | | | x |
| **Aves** | | | | | | | | | | | | | | | | |
| Accipitridae | | | | | | | | | | | | | | | | |
| *Accipiter nisus* | VU | 2 | 0.0 | 0.3 | 0.1 | | | | x | | | x | | | | |
| Accipitridae indet, | | 2 | 0.0 | 0.2 | 0.1 | | | | | | x | | | | | |
| *Buteo buteo* | | 50 | 0.6 | 0.5 | 0.5 | 54 | 0.1 | 0.0 | x | | x | | x | | | x |
| Alcedinidae | | | | | | | | | | | | | | | | |
| *Alcedo atthis* | VU | 3 | 0.0 | 0.2 | 0.1 | | | | | | | x | | | | |
| Anatidae | | | | | | | | | | | | | | | | |
| *Anas acuta* | EN | | | | | 16877 | 0.4 | 0.1 | | x | x | | | x | | |
| *Anas clypeata* | VU | | | | | 1321 | 0.4 | 0.1 | | | | | x | x | x | |
| *Anas crecca* | VU | 3 | 0.0 | 0.2 | 0.1 | 17397 | 0.9 | 0.3 | | x | x | x | x | x | x | |
| *Anas platyrhynchos* | | 1422 | 16.2 | 1.0 | 0.9 | 816794 | 1.0 | 0.9 | x | x | x | x | x | x | x | x |
| *Anatinae indet,* | | | | | | 239 | 0.1 | 0.0 | | | | | x | | | |
| *Anser albifrons* | NT | | | | | 3844 | 0.6 | 0.2 | | x | | | | x | x | x |

| | | | | | | | | | | | | | | | | |
|---|---|---|---|---|---|---|---|---|---|---|---|---|---|---|---|---|
| *Anser anser* | N2000 | 45 | 0.5 | 0.3 | 0.3 | 1989848 | 0.9 | 0.7 | | | x | x | x | x | x | x |
| *Anser indet,* | | | | | | 111966 | 1.0 | 0.6 | | x | x | x | x | x | x | x |
| *Aythya ferina* | VU | | | | | 7481 | 0.4 | 0.1 | | x | | | | x | x | |
| *Aythya fuligula* | NT | | | | | 52391 | 0.9 | 0.3 | | x | x | x | | x | x | x |
| *Branta canadensis* | | | | | | 327033 | 0.7 | 0.3 | | x | | x | x | x | x | |
| *Bucephala clangula* | VU | 9 | 0.1 | 0.3 | 0.3 | | | | | | x | x | | | | |
| *Cairina moscata* | DOM | 1 | 0.0 | 0.2 | 0.1 | 4968 | 0.1 | 0.0 | x | x | | | | | | |
| *Cygnus cygnus* | N2000 | | | | | 34090 | 0.4 | 0.2 | | | | | | x | x | x |
| *Cygnus olor* | | | | | | 4899 | 0.4 | 0.1 | | x | | | | x | | x |
| *Mareca penelope* | CR | | | | | 388197 | 0.9 | 0.5 | | x | x | x | x | x | x | |
| *Mergellus albellus* | | | | | | 18604 | 0.3 | 0.1 | | | | | | x | x | |
| *Mergus merganser* | | 1 | 0.0 | 0.2 | 0.1 | 130 | 0.1 | 0.0 | | x | x | | | | | |
| *Tadorna tadorna* | | | | | | 91 | 0.1 | 0.0 | | x | | | | | | |
| Ardeidae | | | | | | | | | | | | | | | | |
| *Botaurus stellaris* | VU, N2000 | 2 | 0.0 | 0.3 | 0.1 | | | | | | x | | x | | | |
| *Ardea cinerea* | | 331 | 3.8 | 0.9 | 0.8 | 59 | 0.1 | 0.1 | x | x | x | x | x | | | x |
| Certhiidae | | | | | | | | | | | | | | | | |
| *Certhia brachydactyla* | | 1 | 0.0 | 0.2 | 0.1 | | | | | | | | x | | | |
| Cinclidae | | | | | | | | | | | | | | | | |
| *Cinclus cinclus* | CR | 4 | 0.1 | 0.3 | 0.2 | | | | x | | x | | | | | |
| Columbidae | | | | | | | | | | | | | | | | |
| *Columba palumbus* | | 642 | 7.3 | 0.9 | 0.8 | 993 | 0.4 | 0.1 | x | | x | x | x | x | | x |
| Corvidae | | | | | | | | | | | | | | | | |
| *Corvidae indet,* | | | | | | 2867 | 0.7 | 0.1 | | x | x | x | | | x | x |
| *Corvus corax* | | | | | | 255 | 0.3 | 0.1 | | x | | | | | x | |
| *Corvus cornix* | | 92 | 1.1 | 0.2 | 0.5 | | | | | | x | | | | | x |
| *Corvus corone* | | | | | | 2873 | 0.7 | 0.2 | | | x | x | x | | x | x |
| *Corvus frugilegus* | | | | | | 4021 | 0.3 | 0.1 | | x | | | | | x | |
| *Garrulus glandarius* | | 44 | 0.5 | 0.3 | 0.4 | | | | x | | x | | | | | |
| Emberizidae | | | | | | | | | | | | | | | | |
| *Emberiza citrinella* | VU | 1 | 0.0 | 0.2 | 0.1 | | | | | | | | | | | x |
| Fringillidae | | | | | | | | | | | | | | | | |
| *Chloris chloris* | NT | 4 | 0.1 | 0.2 | 0.1 | | | | | | x | | | | | |
| *Coccothraustes coccothraustes* | | 7 | 0.1 | 0.2 | 0.1 | | | | | | x | | | | | |

| | | | | | | | | | | | | | | | | |
|---|---|---|---|---|---|---|---|---|---|---|---|---|---|---|---|---|
| *Fringilla coelebs* | | 27 | 0.3 | 0.2 | 0.3 | 230 | 0.4 | 0.1 | | | x | | | x | | x |
| *Loxia curvirostra* | | 9 | 0.1 | 0.2 | 0.1 | | | | | | x | | | | | |
| *Spinus spinus* | NT | 1 | 0.0 | 0.2 | 0.1 | | | | | | | | x | | | |
| Laridae | | | | | | | | | | | | | | | | |
| *Larinae indet,* | | | | | | 19035 | 0.3 | 0.1 | | | | | x | | x | |
| Motacillidae | | | | | | | | | | | | | | | | |
| *Anthus pratensis* | | 1 | 0.0 | 0.2 | 0.1 | | | | | | x | | | | | |
| *Motacilla alba* | | 92 | 1.1 | 0.5 | 0.4 | | | | | | x | x | | | | x |
| *Motacilla cinerea* | VU | 26 | 0.3 | 0.5 | 0.3 | | | | x | x | x | | | | | |
| *Motacilla flava* | | 1 | 0.0 | 0.2 | 0.1 | | | | x | | | | | | | |
| Motacilla indet, | | 2 | 0.0 | 0.2 | 0.1 | | | | | | x | | | | | |
| Muscicapidae | | | | | | | | | | | | | | | | |
| *Erithacus rubecula* | | 121 | 1.4 | 0.7 | 0.6 | | | | x | | x | | x | | | x |
| *Phoenicurus phoenicurus* | | 3 | 0.0 | 0.3 | 0.1 | | | | x | | x | | | | | |
| Paridae | | | | | | | | | | | | | | | | |
| *Cyanistes caeruleus* | | 3 | 0.0 | 0.2 | 0.2 | | | | | | x | | | | | |
| *Parus major* | | 13 | 0.2 | 0.5 | 0.3 | | | | x | | x | | x | | | |
| *Periparus ater* | | 1 | 0.0 | 0.2 | 0.1 | | | | | | x | | | | | |
| Passeridae | | | | | | | | | | | | | | | | |
| *Passer montanus* | | 4 | 0.1 | 0.2 | 0.1 | | | | | | x | | | | | |
| Phalacrocoracidae | | | | | | | | | | | | | | | | |
| *Phalacrocorax carbo* | | 3 | 0.0 | 0.3 | 0.2 | 33732 | 0.6 | 0.2 | | | x | | x | x | x | x |
| Phasianidae | | | | | | | | | | | | | | | | |
| *Gallus gallus* | DOM | 2 | 0.0 | 0.2 | 0.1 | 376 | 0.3 | 0.1 | x | | | | | x | | x |
| *Phasianus colchicus* | | 225 | 2.6 | 0.5 | 0.6 | 3545 | 0.6 | 0.1 | x | | x | x | x | | | x |
| *Meleagris gallopavo* | DOM | | | | | 801 | 0.1 | 0.1 | | | x | | | | | |
| Phylloscopidae | | | | | | | | | | | | | | | | |
| *Phylloscopus trochilus* | VU | 1 | 0.0 | 0.2 | 0.1 | | | | x | | | | | | | |
| Picidae | | | | | | | | | | | | | | | | |
| *Dendrocopos major* | | 5 | 0.1 | 0.2 | 0.2 | | | | | | x | | | | | |
| Podicipedidae | | | | | | | | | | | | | | | | |
| *Podiceps cristatus* | | | | | | 3703 | 0.1 | 0.1 | | | | | | | x | |
| *Podiceps grisegena* | | 1 | 0.0 | 0.2 | 0.1 | | | | | | x | | | | | |
| *Tachybaptus ruficollis* | | 13 | 0.2 | 0.2 | 0.1 | 13444 | 0.7 | 0.3 | | | x | x | x | x | x | |

| | | | | | | | | | | | | | | | |
|---|---|---|---|---|---|---|---|---|---|---|---|---|---|---|---|
| Rallidae | | | | | | | | | | | | | | | |
| *Fulica atra* | VU | 71 | 0.8 | 0.5 | 0.4 | 46088 | 1.0 | 0.5 | | x | x | x | x | x | x | x |
| *Gallinula chloropus* | VU | 65 | 0.7 | 0.9 | 0.3 | 177 | 0.3 | 0.1 | x | x | x | x | x | | x | x |
| *Rallus aquaticus* | | 7 | 0.1 | 0.2 | 0.1 | | | | | | x | | | | | |
| Scolopacidae | | | | | | | | | | | | | | | |
| *Tringa ochropus* | *EN* | 3 | 0.0 | 0.3 | 0.1 | | | | x | | x | | | | | |
| *Scolopax rusticola* | | 12 | 0.1 | 0.5 | 0.2 | 954 | 0.3 | 0.1 | x | | x | x | | x | | x |
| Sittidae | | | | | | | | | | | | | | | |
| *Sitta europaea* | | 9 | 0.1 | 0.2 | 0.2 | | | | | | x | | | | | |
| Strigidae | | | | | | | | | | | | | | | |
| *Strix aluco* | | 6 | 0.1 | 0.3 | 0.2 | | | | | | x | x | | | | |
| Sturnidae | | | | | | | | | | | | | | | |
| *Sturnus vulgaris* | VU | 4 | 0.1 | 0.2 | 0.1 | | | | | | x | | | | | |
| Sylviidae | | | | | | | | | | | | | | | |
| *Sylvia atricapilla* | | 2 | 0.0 | 0.3 | 0.1 | | | | | | x | | x | | | |
| *Sylvia communis* | | 1 | 0.0 | 0.2 | 0.1 | | | | | | x | | | | | |
| Troglodytidae | | | | | | | | | | | | | | | |
| *Troglodytes troglodytes* | | 24 | 0.3 | 0.3 | 0.3 | | | | | | x | | | | | x |
| Turdidae | | | | | | | | | | | | | | | |
| *Turdus iliacus* | | 7 | 0.1 | 0.3 | 0.2 | | | | x | | x | | | | | |
| *Turdus* indet, | | 178 | 2.0 | 0.7 | 0.8 | 9921 | 0.7 | 0.2 | x | x | x | x | x | | x | x |
| *Turdus merula* | | 710 | 8.1 | 0.7 | 0.9 | | | | x | | x | | x | | | x |
| *Turdus philomelos* | | 119 | 1.4 | 0.5 | 0.6 | 7880 | 0.4 | 0.1 | | x | x | x | | x | | x |
| *Turdus pilaris* | | 1 | 0.0 | 0.2 | 0.1 | | | | | x | | | | | | |
| **Mammalia** | | | | | | | | | | | | | | | |
| Bovidae | | | | | | | | | | | | | | | |
| *Bos taurus* | DOM | 2 | 0.0 | 0.2 | 0.1 | 173318 | 1.0 | 0.8 | | x | x | x | x | x | x | x |
| *Ovis aries* | DOM | | | | | 89 | 0.1 | 0.0 | | | x | | | | | |
| Canidae | | | | | | | | | | | | | | | |
| *Canis lupus* | DOM | 17 | 0.2 | 0.5 | 0.3 | 20491 | 1.0 | 0.3 | | x | x | x | x | x | x | x |
| *Vulpes vulpes* | NT | 532 | 6.1 | 0.8 | 0.7 | 118858 | 0.7 | 0.2 | x | x | x | x | x | x | x | x |
| Cervidae | | | | | | | | | | | | | | | |
| *Capreolus capreolus* | | 1172 | 13.4 | 0.5 | 0.7 | 95391 | 0.7 | 0.4 | x | | x | x | x | | x | x |
| Cervidae indet, | | 3 | 0.0 | 0.2 | 0.1 | | | | | | x | | | | | |

| | | | | | | | | | | | | | | | |
|---|---|---|---|---|---|---|---|---|---|---|---|---|---|---|---|
| *Cervus elaphus* | | 52 | 0.6 | 0.2 | 0.4 | 17870 | 0.7 | 0.3 | | | x | x | x | x | x | |
| *Dama dama* | | 3 | 0.0 | 0.2 | 0.1 | | | | | | x | | | | | |
| Cricetidae | | | | | | | | | | | | | | | | |
| *Arvicola amphibious* | | 58 | 0.7 | 1.0 | 0.4 | | | | x | x | x | x | x | | | x |
| Arvicolinae indet, | | 4 | 0.1 | 0.5 | 0.2 | 9889 | 1.0 | 0.5 | x | x | x | x | x | x | x | x |
| *Microtus agrestis* | | 2 | 0.0 | 0.3 | 0.1 | | | | x | | | x | | | | |
| *Myodes glareolus* | | 1 | 0.0 | 0.2 | 0.1 | 43105 | 0.9 | 0.3 | | x | x | x | x | x | | x |
| Equidae | | | | | | | | | | | | | | | | |
| *Equus ferus* | DOM | | | | | 233 | 0.1 | 0.1 | | | x | | | | | |
| Erinaceidae | | | | | | | | | | | | | | | | |
| *Erinaceus europaeus* | | 1 | 0.0 | 0.2 | 0.1 | | | | | | x | | | | | |
| Felidae | | | | | | | | | | | | | | | | |
| *Felis catus* | DOM | 7 | 0.1 | 0.3 | 0.2 | 16582 | 0.1 | 0.1 | | | x | x | x | | | |
| Leporidae | | | | | | | | | | | | | | | | |
| *Lepus europaeus* | | 3 | 0.0 | 0.2 | 0.1 | | | | | | x | | | | | |
| Muridae | | | | | | | | | | | | | | | | |
| *Apodemus flavicollis* | | | | | | 1302 | 0.6 | 0.1 | | | | x | x | x | x | |
| *Micromys minutus* | | | | | | 638 | 0.3 | 0.1 | | | | x | x | | | |
| Muridae indet, | | 73 | 0.8 | 1.0 | 0.7 | | | | x | x | x | x | x | | | x |
| *Rattus norvegicus* | | 719 | 8.2 | 1.0 | 1.0 | 20456 | 0.9 | 0.3 | x | x | x | x | x | x | | x |
| Mustelidae | | | | | | | | | | | | | | | | |
| *Lutra lutra* | VU, N2000 | 35 | 0.4 | 0.3 | 0.3 | | | | | | x | x | | | | |
| *Martes martes* | NT, N2000 | 572 | 6.5 | 0.9 | 0.8 | | | | x | | x | x | x | | | x |
| *Meles meles* | | 17 | 0.2 | 0.2 | 0.3 | | | | | | x | | | | | |
| *Mustela erminea* | NT | 15 | 0.2 | 0.7 | 0.3 | | | | | x | x | x | | | | x |
| *Mustela nivalis* | NT | 2 | 0.0 | 0.3 | 0.1 | | | | | | x | x | | | | |
| *Mustela putorius* | NT | 76 | 0.9 | 0.7 | 0.7 | | | | x | x | x | | x | | | |
| Mustelidae indet, | | 76 | 0.9 | 0.8 | 0.8 | | | | x | | x | x | x | | | x |
| *Neovison vison* | | 342 | 3.9 | 0.9 | 0.8 | | | | x | | x | x | x | | | x |
| Sciuridae | | | | | | | | | | | | | | | | |
| *Sciurus vulgaris* | | 234 | 2.7 | 0.3 | 0.3 | 91 | 0.1 | 0.0 | x | | x | | | | | |
| Soricidae | | | | | | | | | | | | | | | | |
| *Neomys fodiens* | NT | 14 | 0.2 | 0.7 | 0.3 | 1413 | 0.1 | 0.1 | x | | | x | x | | | x |
| *Sorex araneus* | | | | | | 1692 | 0.4 | 0.1 | | | | | x | x | | x |

| | | | | | | | | | | | | | | | | |
|---|---|---|---|---|---|---|---|---|---|---|---|---|---|---|---|---|
| *Sorex minutus* | | | | | | 339 | 0.1 | 0.0 | | | x | | | | | |
| Soricidae indet, | | 9 | 0.1 | 0.7 | 0.3 | | | | | x | x | x | | | | x |
| *Myomorpha* indet, | | 191 | 2.2 | 0.8 | 0.7 | | | | x | | x | x | x | | | x |
| Suidae | | | | | | | | | | | | | | | | |
| *Sus scrofa* | DOM | | | | | 141520 | 1.0 | 0.7 | | x | x | x | x | x | x | x |
| **Squamata** | | | | | | | | | | | | | | | | |
| Colubridae | | | | | | | | | | | | | | | | |
| *Natrix natrix* | | 2 | 0.0 | 0.2 | 0.1 | | | | | | | x | | | | |
| **Total** | | **8674** | | | | **12166198** | | | **36** | **51** | **104** | **66** | **67** | **49** | **49** | **60** |

**Table S10.** Corrected taxa from eDNA samples. Bacteria and other procaryotes were found but discarded completely. All vertebrates detected by eDNA are explained in this table. The column named 'Action' indicates what evaluation was done given the species inferred in the left most column named 'Given species'. The prefiltering script was applied after sequences had been compared with the NCBI GenBank database using BLAST, but before stacked bar plots were prepared. Notes on species from Månsson (2018) samples from sequencing libraries M-R.

| Given species | Action | Reasoning for action |
|---|---|---|
| *Ablennes hians* | Removed | Contamination only found in Månssons samples (2018) (Library M-R). Fish from Qatar |
| *Acanthurus olivaceus* | Removed | Prefiltering script connected it with *Gasterosteus aculeatus*, but only 85 % identical, thereby disregarded. Only found in mock |
| *Alces alces* | *Cervus elaphus* | Corrected for via prefiltering script |
| *Amaurornis phoenicurus* | *Gallinula chloropus* | Corrected for via prefiltering script |
| *Anampses neoguinaicus* | *Anampses twistti* | Closely related genetically, only found in mock samples |
| *Anas chathamica* | *Anas platyrhynchos* | Corrected for via prefiltering script |
| *Anguilla dieffenbachii* | *Anguilla anguilla* | Corrected for via prefiltering script |
| *Anser caerulenscens* | *Anser* indet. | Genetically identical to more than one *Anser* species that are possible in the area |
| *Anser cygnoides* | *Anser anser* | Swan goose has not been spotted in the area for 10 years and is extremely rare in DK. Similar genetics in the 12S gene to the Greylag goose, which is very frequent in the area |
| *Baleana mysticetus* | Removed | Contamination only found in Månssons samples (Library M-R). |
| *Bodianus anthiodes* | *Bodianus axillaris* | Closely related genetically, only found in mock samples |
| *Branta sandvicensis* | *Branta canadensis* | Corrected for via prefiltering script |
| *Cephalophus natalensis* | Removed | Few reads and only in the mock samples. |
| *Chaerodon selene* | Removed | Lab contaminant, only found in mock, under five reads |
| *Chaetodon rafflesi* | Removed | Lab contaminant, only found in mock, under five reads |
| *Chanos chanos* | Removed | Contamination only found in Månssons samples (Library M-R). Fish from Qatar |
| *Clupea pallasii* | Removed | Lab contaminant only found in mock. Under five reads |
| *Corvus brachyrhynchos* | *Corvidae* indet. | Only three NCBI matches for this sequence. One is a subspecies of *Corvus corone* |
| *Corvus hawaiiensis* | *Corvus corone* | Corrected for via prefiltering script |
| *Corvus splendens* | *Corvus corax* | Genetic match on NCBI GenBank |

**Table S11.** Species list of all the vertebrates found in Åmosen, by the means of literature (traditional surveys). Summary of presence of vertebrate taxa fauna found in Nature Park Åmosen through the last ca. 20 years. Data from Atlas over Danske Ferskvandsfisk (Danish Freshwater Fish Atlas) (Carl & Møller 2012) and additional information from the updated atlas database (provided by Henrik Carl and Peter Rask Møller); Dansk Pattedyratlas (Danish Mammal Atlas) (Baagøe & Jensen 2007); Fuglenes Danmark (The birds of Denmark) (Grell 1998) and data from Danish Ornithology Society (DOF) provided by Michael Fink. The national biodiversity portal Arter.dk was also consulted. (†) Presumed locally extinct or not returned. (I) only found in a little spot in the outer edges of Åmosen. (II) Found at 'Flasken' (a border between the stream and the ocean. According to the Danish IUCN Red List: (CR) critically endangered, (EN) endangered, (VU) vulnerable, (NT) near threatened, these statuses is made from the 2019 updated list, note that there can be a difference if the birds are migratory or breeding in the area. The comments also include whether the species is Natura 2000 protected in the area and what directive annex they are protected in if they are (European Commission 2009, Naturstyrelsen 2012).

| Family | Species | Common name | Outlier | Comment | eDNA | CT |
|---|---|---|---|---|---|---|
| **Agnatha** | | | | | | |
| Petromyzontidae | *Lampetra fluviatilis†* | European river lamprey | Yes | Annex II+V, Natura 2000, latest obs. early 1900s | | |
| | *Lampetra planeri†* | Brook lamprey | Yes | Annex II, Natura 2000, latest obs. early 1900s | | |
| **Actinopterygii** | | | | | | |
| Anguillidae | ***Anguilla anguilla*** | European eel | | CR | X | |
| Cobitidae | ***Cobitis taenia*** | Spined loach | | Annex II, Natura 2000 | X | |
| Cyprinidae | ***Abramis brama*** | Freshwater bream | | | X | |
| | ***Alburnus alburnus*** | Common bleak | | | X | |
| | ***Carassius auratus*** | Goldfish | | Non-indigoens | X | |
| | *Carassius carassius* | Crucian carp | | | | |

| | | | | | |
|---|---|---|---|---|---|
| | *Ctenopharyngodon idella* | Grass carp | | Non-indigoens | |
| | ***Cyprinus carpio*** | Common carp | | | X |
| | ***Gobio gobio*** | Gudgeon | | | X |
| | ***Leucaspius delineatus*** | Sunbleak | | | X |
| | ***Leuciscus idus*** | Ide | | | X |
| | ***Rutilus rutilus*** | Common roach | | | X |
| | ***Scardinius erythrophthalmus*** | Common rudd | | | X |
| | ***Tinca tinca*** | Tench | | | X |
| Esocidae | ***Esox lucius*** | Northern pike | | | X |
| Gasterosteidae | ***Gasterosteus aculeatus*** | Three-spined stickleback | | | X |
| | ***Pungitius pungitius*** | Nine-spined stickleback | | | X |
| Lotidae | *Lota lota*† | Burbot | Yes | Latest obs. 1927 | |
| Percidae | ***Gymnocephalus cernua*** | Ruffe | | | X |
| | ***Perca fluviatilis*** | European perch | | | X |
| | ***Sander lucioperca*** | Zander | | | X |
| Pleuronectidae | *Platichthys flesus* | European flounder | | | |
| Salmonidae | ***Oncorhynchus mykiss*** | Rainbow trout | | Non-indigoens | X |
| | ***Salmo trutta*** | Brown trout | | | X |
| Siluridae | *Silurus glanis* | Wels catfish | | | |
| **Amphibia** | | | | | |
| Bufonidae | ***Bufo bufo*** | Common toad | | | X |

| | | | | | | |
|---|---|---|---|---|---|---|
| | *Epidalea calamitaII* | Natterjack toad | Yes | EN, Natura 2000, Annex IV | | |
| Ranidae | *Pelophylax esculentus* | Edible frog | | Natura 2000, Annex V | | |
| | *Rana arvalis* | Moor frog | | Natura 2000, Annex IV | | |
| | ***Rana temporaria*** | Common frog | | NT | X | |
| Salamandridae | ***Lissotriton vulgaris*** | Smooth newt | | | X | |
| | ***Triturus cristatus*** | Northern crested newt | | Natura 2000, Annex II+IV | X | |
| **Squamata** | | | | | | |
| Colubridae | ***Natrix natrix*** | Grass snake | | | | X |
| Viperidae | *Vipera berus* | European viper | | | | |
| Anguidae | *Anguis fragilis* | Slowworm | | | | |
| Lacertidae | *Lacerta agilis* | Sand lizard | | VU, Natura 2000, Annex IV | | |
| | *Zootoca vivipara* | Vivipaous lizard | | | | |
| **Mammalia** | | | | | | |
| Canidae | ***Vulpes vulpes*** | Red fox | | NT | X | X |
| Cervidae | ***Capreolus capreolus*** | Roe deer | | | X | X |
| | ***Cervus elaphus*** | Red deer | | | X | X |
| | *Cervus nippon* | Sika deer | | Non-indigoens | | |
| | ***Dama dama*** | Fallow deer | | Non-indigoens | | X |
| Cricetidae | ***Arvicola amphibious*** | European water vole | | | | X |
| | ***Microtus agrestis*** | Field vole | | | | X |
| | ***Myodes glareolus*** | Bank vole | | | X | X |

| | | | | | | |
|---|---|---|---|---|---|---|
| Erinaceidae | ***Erinaceus europaeus*** | European hedgehog | | | | X |
| Leporidae | ***Lepus europaeus*** | European hare | | | | X |
| | *Oryctolagus cuniculusI* | European rabbit | Yes | Relatively rare | | |
| Muridae | ***Apodemus flavicollis*** | Yellow-necked mouse | | | X | |
| | *Apodemus sylvaticus* | Wood mouse | | NT | | |
| | ***Micromys minutus*** | Eurasian harvest mouse | | | X | |
| | *Mus musculus* | House mouse | | NT | | |
| | ***Rattus norvegicus*** | Brown rat | | Invasive | X | X |
| Mustelidae | ***Lutra lutra*** | Eurasian otter | | VU, Natura 2000, Annex II+IV, relatively rare | | X |
| | *Martes foina* | Beech marten | | NT | | |
| | ***Martes martes*** | Pine marten | | NT, Natura 2000, Annex V | | X |
| | ***Meles meles*** | European badger | | | | X |
| | ***Mustela erminea*** | Stoat | | NT | | X |
| | ***Mustela nivalis*** | Least weasel | | NT, relatively rare | | X |
| | ***Mustela putorius*** | Polecat | | NT, Annex V | | X |
| | ***Neovison vison*** | American mink | | Invasive species | | X |
| Phocidae | *Phoca vitulina*[II] | Harbour seal | Yes | Natura 2000, Annex II+V | | |
| Phocoenidae | *Phocoena phocoena*[II] | Habour porpoise | Yes | Natura 2000, Annex II+IV | | |
| Sciuridae | ***Sciurus vulgaris*** | Red squirrel | | | X | X |
| Soricidae | ***Neomys fodiens*** | Eurasian water | | NT | X | X |

| | | | | | | |
|---|---|---|---|---|---|---|
| | | shrew | | | | |
| | ***Sorex araneus*** | Common shrew | | | X | |
| | ***Sorex minutus*** | Eurasian pygmy shrew | | | X | |
| Talpidae | *Talpa europaea* | European mole | | | | |
| Vespertilionidae | *Eptesicus serotinus* | Serotine bat | | Annex IV | | |
| | *Myotis daubentonii* | Daubenton's bat | | Annex IV | | |
| | *Nyctalus noctula* | Common noctule | | Annex IV | | |
| | *Pipistrellus nathusii* | Nathusius' pipistrelle | | Annex IV | | |
| | *Pipistrellus pygmaeus* | Soprano pipistrelle | | Annex IV | | |
| | *Plecotus auritus* | Brown long-eared bat | | Annex IV | | |
| **Aves** | | | | | | |
| Accipitridae | *Accipiter gentilis* | Northern goshawk | | VU, Annex I | | |
| | ***Accipiter nisus*** | Eurasian sparrowhawk | | VU, Annex I | | X |
| | *Aquila chrysaetos* | Golden eagle | | CR, Annex I, relatively rare | | |
| | ***Buteo buteo*** | Common buzzard | | | X | X |
| | *Buteo lagopus* | Rough-legged buzzard | | | | |
| | *Circus aeruginosus* | Western-marsh harrier | | Natura 2000, Annex I | | |

| | | | | | | |
|---|---|---|---|---|---|---|
| | *Circus cyaneus* | Hen harrier | | Annex I | | |
| | *Haliaeetus albicilla* | White-tailed eagle | | NT, Natura 2000, Annex I | | |
| | *Milvus milvus* | Red kite | | VU, Natura 2000, Annex I | | |
| | *Pernis apivorus* | European honey buzzard | | NT, Annex I | | |
| Acrocephalidae | *Acrocephalus arundinaceus* | Great reed warbler | Yes | CR, relatively rare | | |
| | *Acrocephalus palustris* | Marsh warbler | | | | |
| | *Acrocephalus schoenobaenus* | Sedge warbler | | | | |
| | *Acrocephalus scirpaceus* | Eurasian reed warbler | | NT | | |
| | *Hippolais icterina* | Icterine warbler | | VU | | |
| Aegithalidae | *Aegithalos caudatus* | Long-tailed tit | | | | |
| Alaudidae | *Alauda arvensis* | Eurasian skylark | | NT, Annex II | | |
| | *Lullula arborea* | Woodlark | Yes | NT, Annex I | | |
| Alcedinidae | *Alcedo atthis* | Common kingfisher | | VU, Annex I | | |
| Alcidae | *Cepphus grylle*[ii] | Black guillemot | Yes | | | |
| | ***Anas acuta*** | Pintail | | EN, Annex II | X | |
| | ***Anas clypeata*** | Northern shoveler | | VU, Annex II | X | |
| | ***Anas crecca*** | Eurasian teal | | VU, Annex II | X | X |
| | ***Anas platyrhynchos*** | Mallard | | Annex II | X | X |
| | *Anas querquedula* | Garganey | | | | |

| | | | | | | |
|---|---|---|---|---|---|---|
| | *Anas strepera* | Gadwall | | | | |
| | ***Anser anser*** | Greylag goose | | Natura 2000, Annex II | X | X |
| | ***Anser albifrons*** | Greater white footed goose | | NT, Natura 2000, Annex II | X | |
| | *Anser brachyrhynchus* | Pink-footed goose | | | | |
| | *Anser fabalis* | Bean goose | | | | |
| | ***Aythya ferina*** | Common pochard | | VU | X | |
| | ***Aythya fuligula*** | Tufted duck | | NT | X | |
| | ***Branta canadensis*** | Canada goose | | Annex II | X | |
| | *Branta leucopsis* | Barnacle goose | | Annex I | | |
| | ***Bucephala clangula*** | Common goldeneye | | VU | | X |
| | *Cygnus columbianus* | Tundra swan | | Natura 2000, Relatively rare | | |
| | ***Cygnus cygnus*** | Whooper swan | | Natura 2000, Annex I | X | |
| | ***Cygnus olor*** | Mute swan | | Annex II | X | |
| | ***Mareca penelope*** | Eurasian widgeon | | CR | X | |
| | ***Mergellus albellus*** | Smew | | Annex I | X | |
| | ***Mergus merganser*** | Common merganser | | Annex II | X | X |
| | *Mergus serrator* | Red-breasted merganser | | Annex II | | |
| | *Somateria mollissima* | Common eider | | NT, Annex II | | |
| | ***Tadorna tadorna*** | Common | | | X | |

| | | | | | | |
|---|---|---|---|---|---|---|
| | | shelduck | | | | |
| Apodidae | *Apus apus* | Common swift | | NT | | |
| Ardeidae | ***Ardea cinerea*** | Grey heron | | | X | X |
| | ***Botaurus stellaris*** | Eurasian bittern | | VU, Natura 2000, Annex I | | X |
| Caprimulgidae | *Caprimulgus europaeus* | European nightjar | | NT, Annex I, relatively rare | | |
| Certhiidae | ***Certhia brachydactyla*** | Short-toed treecreeper | | Relatively rare | | X |
| | *Certhia familiaris* | Eurasian treecreeper | | | | |
| Charadriidae | *Calidris alba*[II] | Sanderling | Yes | | | |
| | *Charadrius dubius* | Little ringed plover | | NT | | |
| | *Charadrius hiaticula* | Common ringed plover | | VU | | |
| | *Pluvialis apricaria* | European golden plover | | CR, Annex II | | |
| | *Pluvialis squatarola*[II] | Grey plover | Yes | Annex II | | |
| | *Vanellus vanellus* | Northern lapwing | | VU, Annex II | | |
| Cinclidae | ***Cinclus cinclus*** | White-throated dipper | | CR | | X |
| Ciconiidae | *Ciconia ciconia* | White stork | | CR, Annex I, relatively rare | | |
| Columbidae | *Columba liviaII* | Rock dove | Yes | Unknown spread. Annex II | | |
| | *Columba oenas* | Stock dove | | Annex II | | |
| | ***Columba palumbus*** | Common wood pidgeon | | Annex II | X | X |

| | | | | | | |
|---|---|---|---|---|---|---|
| | *Streptopelia decaocto* | Eurasian collared dove | | NT, Annex II | | |
| | *Streptopelia turtur* | European turtledove | Yes | EN, Annex II, rare | | |
| Corvidae | *Coloeus monedula* | Western jackdaw | | | | |
| | ***Corvus corax*** | Common raven | | | X | |
| | ***Corvus cornix*** | Hooded crow | | | | X |
| | ***Corvus corone*** | Carrion crow | Yes | Annex II | X | |
| | ***Corvus frugilegus*** | Rook | | Annex II | X | |
| | ***Garrulus glandarius*** | Eurasian jay | | Annex II | | X |
| | *Nucifraga caryocatactes* | Northern nutcracker | | Possibly missing in the area, relatively rare | | |
| | *Pica pica* | Eurasian magpie | | Annex II | | |
| Cuculidae | *Cuculus canorus* | Common cuckoo | | NT | | |
| | *Emberiza calandra* | Corn bunting | | NT | | |
| | ***Emberiza citrinella*** | Yellowhammer | | VU | | X |
| Emberizidae | *Emberiza schoeniclus* | Common red bunting | | NT | | |
| Falconidae | *Falco columbarius* | Merlin | | Annex I, | | |
| | *Falco peregrinus* | Peregrine falcon | | VU, Annex I, earlier thought to be missing in the area | | |
| | *Falco subbuteo* | Eurasian hobby | | CR, relatively rare | | |
| | *Falco tinnunculus* | Common kestrel | | | | |
| Fringillidae | *Acanthis flammea* | Common redpoll | | | | |
| | *Carduelis carduelis* | European | | | | |

| | | | | | | |
|---|---|---|---|---|---|---|
| | | goldfinch | | | | |
| | *Carduelis flavirostris* | Twite | | | | |
| | ***Chloris chloris*** | European greenfinch | | NT | | X |
| | *Coccothraustes coccothraustes* | Hawfinch | | | | |
| | ***Fringilla coelebs*** | Common chaffinch | | | X | X |
| | *Fringilla montifringilla* | Brambling | | | | |
| | *Linaria cannabina* | Common linnet | | | | |
| | ***Loxia curvirostra*** | Red crossbill | | | | X |
| | *Pyrrhula pyrrhula* | Eurasian bullfinch | | | | |
| | *Serinus serinus* | European serin | Yes | CR, relatively rare | | |
| | ***Spinus spinus*** | European sisken | | NT | | X |
| Gruidae | *Grus grus* | Common crane | | Annex I | | |
| Haematopodidae | *Haematopus ostralegus* | Eurasian oystercatcher | | Annex II | | |
| Hirundinidae | *Delichon urbicum* | Common house martin | | | | |
| | *Hirundo rustica* | Barn swallow | | | | |
| | *Riparia riparia* | Collared sand martin | | NT | | |
| Laniidae | *Lanius collurio* | Red-backed shrike | | Annex I | | |
| Laridae | *Chroicocephalus ridibundus* | Black-headed gull | | EN | | |
| | *Larus canus* | Mew gull | | Annex II | | |

| | | | | | | |
|---|---|---|---|---|---|---|
| | *Larus fuscus*[II] | Lessor black-backed gull | Yes | Annex II | | |
| | *Larus marinus* | Great black-backed gull | | Annex II | | |
| | *Hydrocoloeus minutus* | Little gull | | CR. Thought to be missing in the area. Relatively rare | | |
| | *Sterna hirundo* | Common tern | | NT, Annex I, Natura 2000 | | |
| | *Sterna paradisaea*[II] | Arctic tern | Yes | VU, Annex I | | |
| | *Sternula albifrons* | Little tern | | VU, Natura 2000 | | |
| | *Thalasseus sandvicensis*[II] | Sandwich tern | Yes | | | |
| | *Locustella fluviatilis* | River warbler | Rare | | | |
| | *Locustella naevia* | Common grasshopper warbler | | | | |
| | *Locustella luscinioides* | Savi's warbler | | CR, relatively rare | | |
| | ***Anthus pratensis*** | Meadow pipit | | | | X |
| | *Anthus trivialis* | Tree pipit | Yes | | | |
| | ***Motacilla alba*** | White wagtail | | | | X |
| | ***Motacilla cinerea*** | Grey wagtail | | VU | | X |
| | ***Motacilla flava*** | Western yellow wagtail | | | | X |
| Muscicapidae | ***Erithacus rubecula*** | European robin | | | | X |
| | *Ficedula hypoleuca* | European pied | | VU | | |

| | | | | | | |
|---|---|---|---|---|---|---|
| | | flycatcher | | | | |
| | *Luscinia luscinia* | Thrush nightingale | | VU | | |
| | *Muscicapa striata* | Spotted flycatcher | | | | |
| | *Oenanthe oenanthe* | Northern wheateater | | VU | | |
| | *Phoenicurus ochruros* | Black redstart | | NT | | |
| | ***Phoenicurus phoenicurus*** | Common redstart | | | | X |
| | *Saxicola rubetra* | Whincat | | | | |
| Oriolidae | *Oriolus oriolus* | Eurasian golden oriole | Yes | CR, Relatively rare | | |
| Pandionidae | *Pandion haliaetus* | Osprey | | CR, Natura 2000, Annex I | | |
| Panuridae | *Panurus biarmicus* | Bearded reedling | | | | |
| Paridae | ***Cyanistes caeruleus*** | Eurasian blue tit | | | | X |
| | ***Parus major*** | Great tit | | | | X |
| | ***Periparus ater*** | Coal tit | | | | X |
| | *Poecile palustris* | Marsh tit | | | | |
| Passeridae | *Passer domesticus* | House sparrow | | | | |
| | ***Passer montanus*** | Eurasian tree sparrow | | | | X |
| Phalacrocoracidae | ***Phalacrocorax carbo*** | Great cormorant | | | X | X |
| Phasianidae | *Perdix perdix* | Grey partridge | | VU, Annex II | | |
| | ***Phasianus colchicus*** | Common pheasant | | Non-indigenous | X | X |

| | | | | | | |
|---|---|---|---|---|---|---|
| Phylloscopidae | *Phylloscopus collybita* | Common chiffchaff | | | | |
| | *Phylloscopus sibilatrix* | Wood warbler | | | | |
| | ***Phylloscopus trochilus*** | Willow warbler | | VU | | X |
| Picidae | ***Dendrocopos major*** | Great spotted woodpecker | | | | X |
| | *Dryocopus martius* | Black woodpecker | Yes | VU, Annex I, relatively rare | | |
| | *Dryobates minor* | Lessor spotted woodpecker | Yes | EN, relatively rare | | |
| Podicipedidae | ***Podiceps cristatus*** | Great crested grebe | | | X | |
| | ***Podiceps grisegena*** | Red-necked grebe | | | | X |
| | *Podiceps nigricollis* | Black-necked grebe | | | | |
| | ***Tachybaptus ruficollis*** | Little grebe | | | X | X |
| Prunellidae | *Prunella modularis* | Dunnock | | | | |
| Rallidae | *Crex crex* | Corncrake | | VU, Annex I, relatively rare | | |
| | ***Fulica atra*** | Eurasian coot | | VU, Annex II | X | X |
| | ***Gallinula chloropus*** | Common moorhen | | VU, Annex II | X | X |
| | *Porzana porzana* | Spotted crake | | EN, Natura 2000, Annex I, relatively rare | | |
| | ***Rallus aquaticus*** | Water rail | | Annex II | | X |
| Recurvirostri | *Recurvirostra* | Pied avocet | | VU, Annex II | | |

| | | | | | | |
|---|---|---|---|---|---|---|
| dae | *avosetta* | | | | | |
| Regulidae | *Regulus ignicapilla* | Common firecrest | | Relatively rare | | |
| | *Regulus regulus* | Goldcrest | | | | |
| Remizidae | *Remiz pendulinus* | Eurasian penduline tit | | CR | | |
| Scolopacidae | *Actitis hypoleucos* | Common sandpiper | | | | |
| | *Calidris alpina* | Dunlin | | EN | | |
| | *Calidris ferruginea* | Curlew sandpiper | | | | |
| | *Calidris pugnax* | Ruff | | EN, Natura 2000 | | |
| | *Gallinago gallinago* | Common snipe | | Annex III | | |
| | *Limosa lapponica* | Bar-tailed godwit | | Annex II | | |
| | *Limosa limosa*[II] | Black-tailed godwit | Yes | VU, Annex II | | |
| | *Lymnocryptes minimus* | Jack snipe | | Annex III, relatively rare | | |
| | *Numenius arquata* | Eurasian curlew | | VU, Annex II | | |
| | *Numenius phaeopus*[II] | Whimbrel | Yes | Annex II | | |
| | ***Scolopax rusticola*** | Eurasian woodcock | | Annex III | X | X |
| | *Tringa glareola* | Wood sandpiper | | EN, Annex I | | |
| | *Tringa nebularia*[II] | Common greenshank | Yes | Annex II | | |
| | ***Tringa ochropus*** | Green sandpiper | | EN | | X |
| | *Tringa totanus* | Common redshank | | NT, Annex II | | |

| | | | | | | |
|---|---|---|---|---|---|---|
| Sittidae | ***Sitta europaea*** | Eurasian nuthatch | | | | X |
| Strigidae | *Asio flammeus* | Short-eared owl | | CR, Annex I, relatively rare | | |
| | *Asio otus* | Long-eared owl | | | | |
| | ***Strix aluco*** | | | | | X |
| Sturnidae | ***Sturnus vulgaris*** | Common starling | | | | X |
| Sylviidae | ***Sylvia atricapilla*** | Eurasian blackcap | | | | X |
| | *Sylvia borin* | Garden warbler | | | | |
| | ***Sylvia communis*** | Common whitethroat | | | | X |
| | *Sylvia curruca* | Lessor whitethroat | | | | |
| Troglodytidae | ***Troglodytes troglodytes*** | Eurasian wren | | | | X |
| Turdidae | ***Turdus iliacus*** | Redwing | | Annex II | | X |
| | ***Turdus merula*** | Blackbird | | Annex II | | X |
| | ***Turdus philomelos*** | Song thrush | | Annex II | X | X |
| | ***Turdus pilaris*** | Fieldfare | | Annex II | | X |
| | *Turdus torquatus* | Ring ouzel | Yes | Relatively rare | | |
| | *Turdus viscivorus* | Mistle thrush | | Annex II | | |

**Supplementary figures**

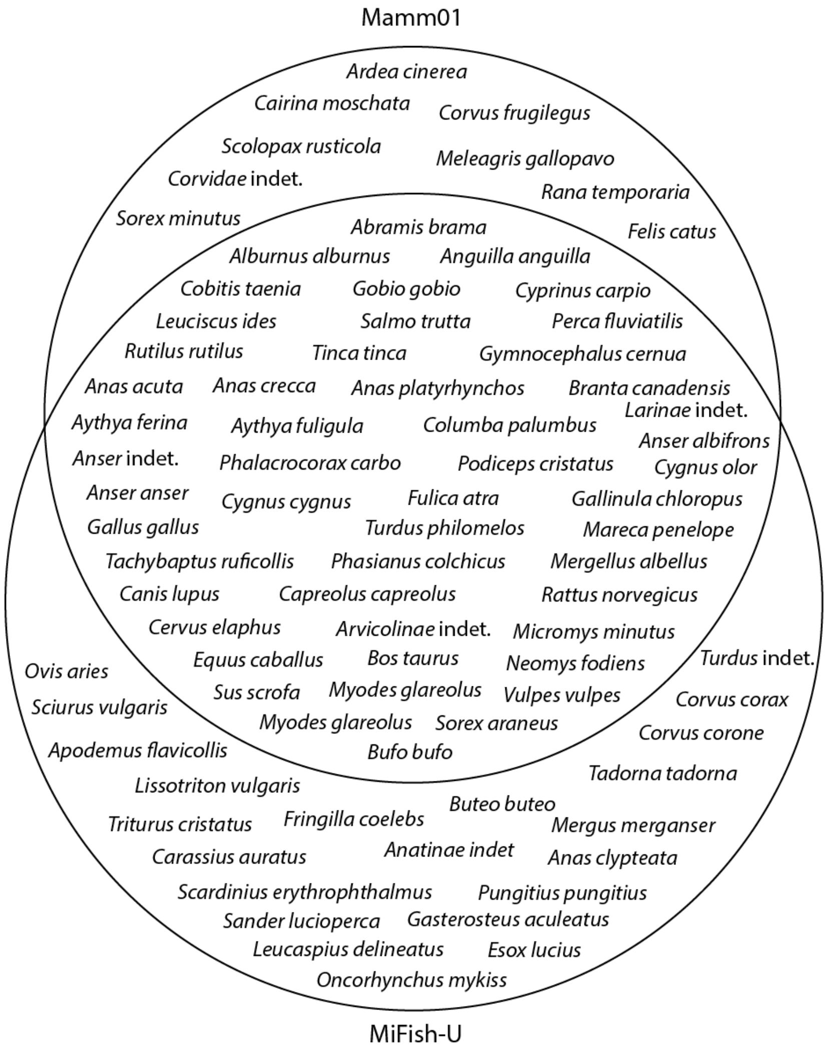

**Supplementary Figure S1. Overview of taxa detected by each primer sets, used in on the water samples collected in Åmosen.** *Venn diagram* showing overlap between the two primer sets: Mamm01 (Taberlet *et al.* 2018a) and MiFish-U (Miya *et al.* 2015). Both primer sets detected 49 taxa, while 10 taxa only were detected with Mamm01 and 23 taxa only detected by MiFish-U.

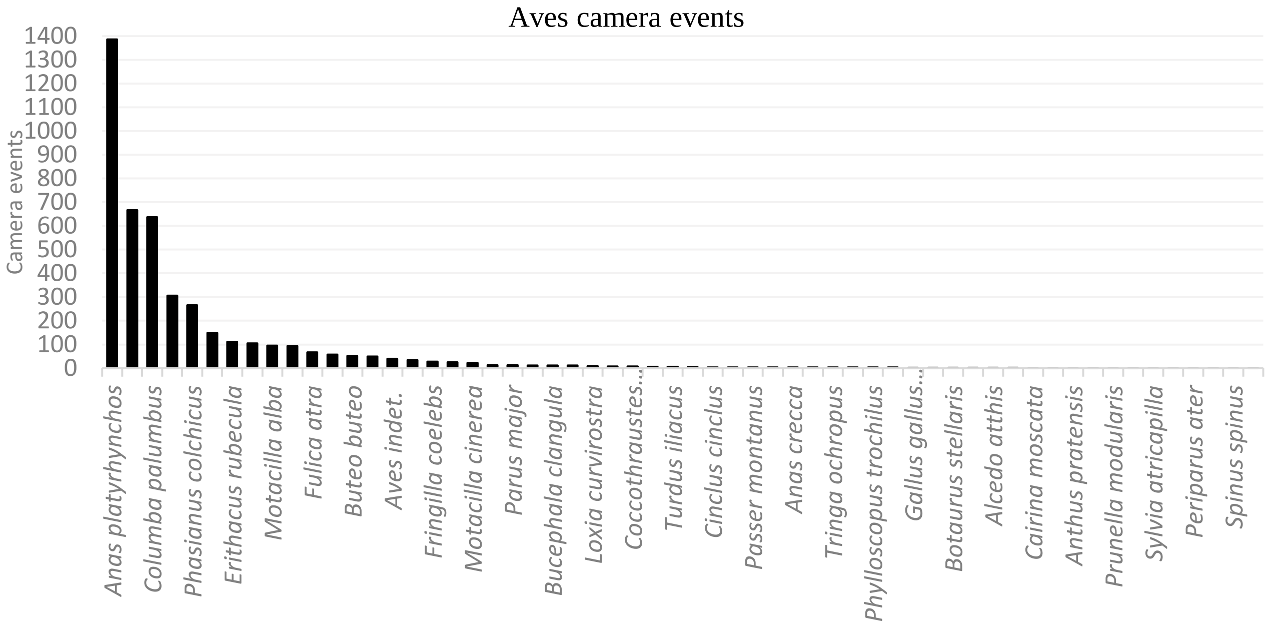

**Supplementary Figure 2.** Camera event (sightings with 30 minutes interval) frequency from all mammal and bird taxa found in Åmosen. A total of 9089 camera events (CE) spread out on 8778 camera days found 87 different taxa. The first graph shows the 29 mammalian taxa found from most frequent to least. The second graph shows the 58 avian taxa, from most frequent to least.

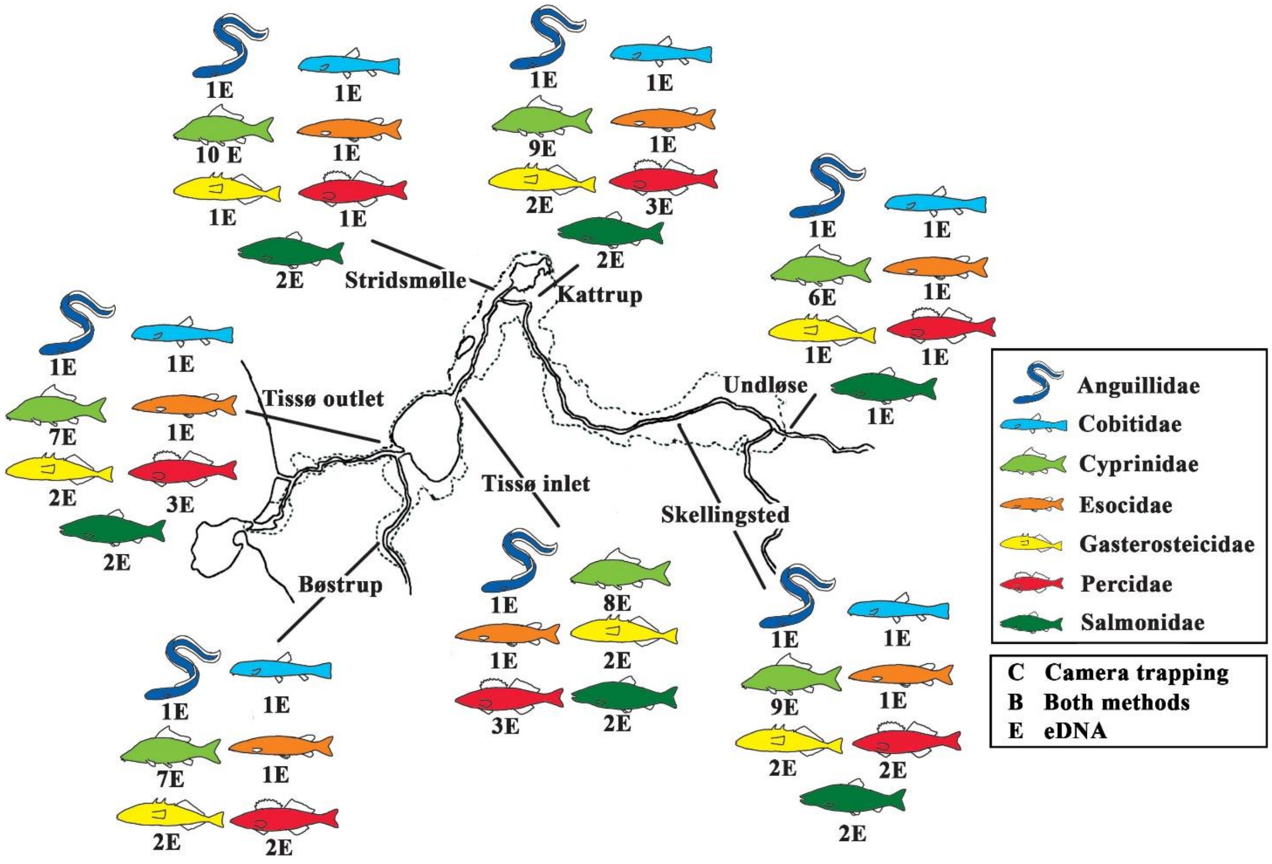

**Suppl. Figure S3. Overview of fish families found in Åmosen.** The qualitative data on fish species found in Åmosen via four years of camera trapping (C), 15 months of eDNA (E), or via both methods (B) sorted into families. Each family has a varying number of maximum detected species in the area: Anguillidae (one species), Cobitidae (one species), Cyprinidae (10 species), Esocidae (one species), Gasterosteicidae (two species), Percidae (three species), and Salmonidae (two species) (see Table 2 for species). Illustrations by AMRH. Animals are not to scale.

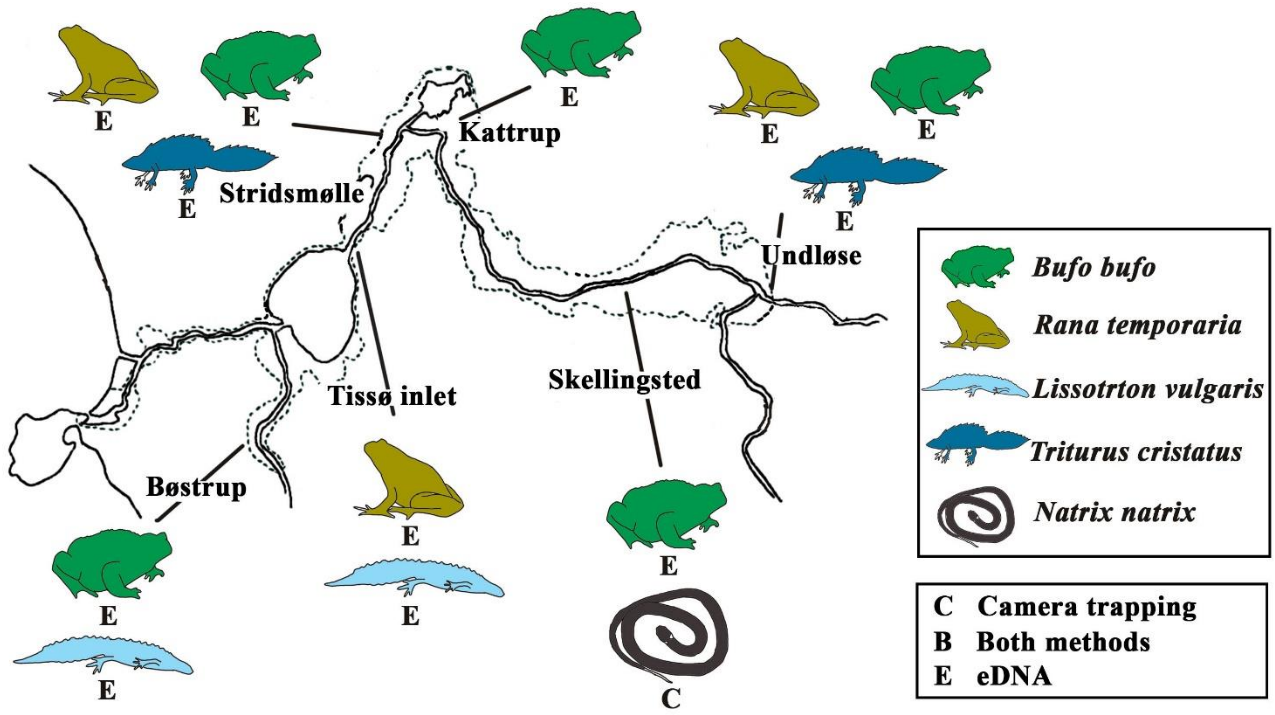

**Suppl. Figure S4. Overview of amphibians and a squamate found in Åmosen.** The qualitative data on the four amphibians, and one squamate detected via four years of camera trapping (C), 15 months eDNA (E), or via both methods (B) (Table 2). Illustrations by AMRH. Animals are not to scale.

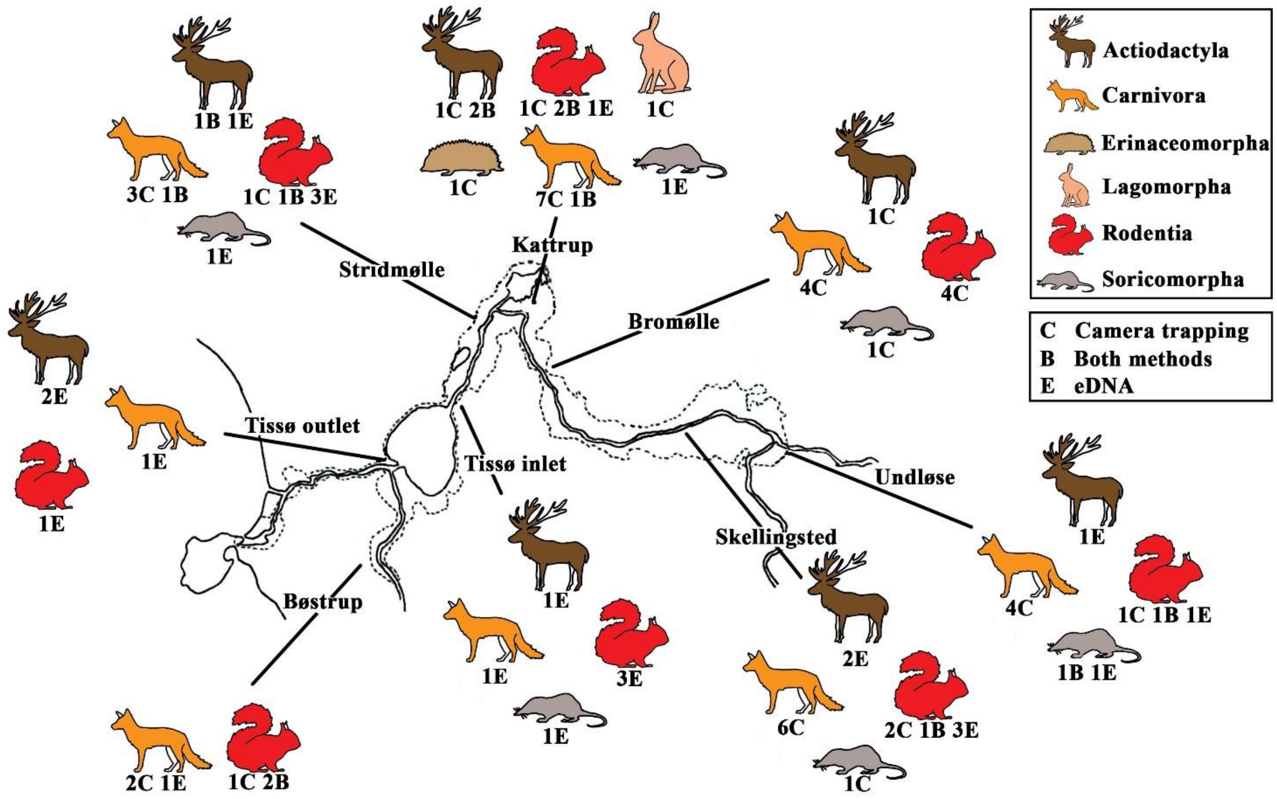

**Suppl. Figure S5. Overview of mammals found in Åmosen**. The qualitative data on the mammal species found in Åmosen via four years of camera trapping (C), 15 months of eDNA (E), or via both methods (B), sorted into orders. Each order has a varying number of maximum detected species in the area: Actiodactyla (three species), Carnivora (eight species), Erinaceomorpha (one species), Lagomorpha (one species), Rodentia (seven species), and Soricomorpha (three species) (see Table 2 for species). Illustrations by AMRH. Animals are not to scale.

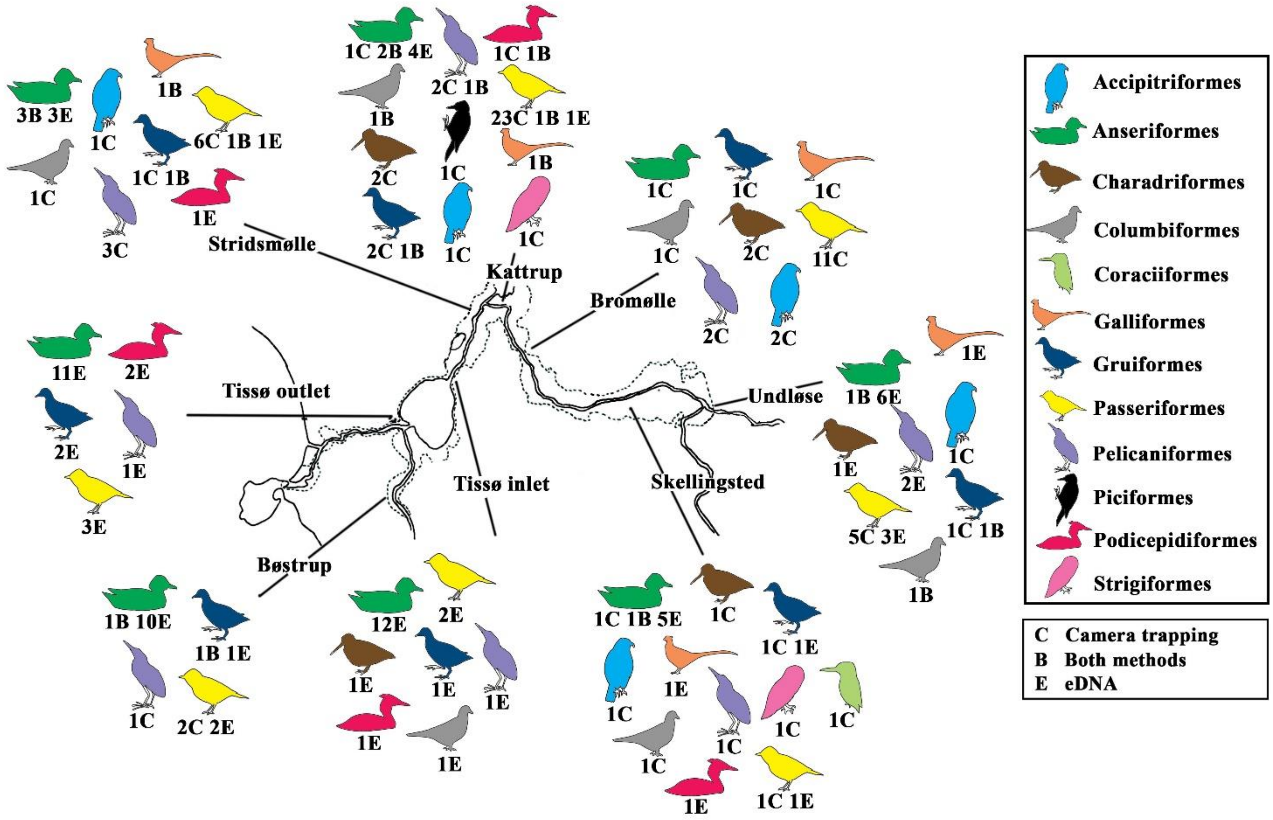

**Suppl. Figure S6 Overview of birds found in Åmosen**. The qualitative data on bird species found in Åmosen via four years of camera trapping (C), 15 months of eDNA (E), or via both methods (B), sorted into orders. Each order has a varying number of maximum detected species in the area: Accipitriformes (two species), Anseriformes (16 species), Charadriiformes (two species), Columbiformes (one species), Coraciiformes (one species), Galliformes (one species), Gruiformes (three species), Passeriformes (33 species), Pelicaniformes (three species), Piciformes (one species), Podicepidiformes (three species), and Strigiformes (one species) (see Table 2 for species). Illustrations by AMRH. Animals are not to scale.

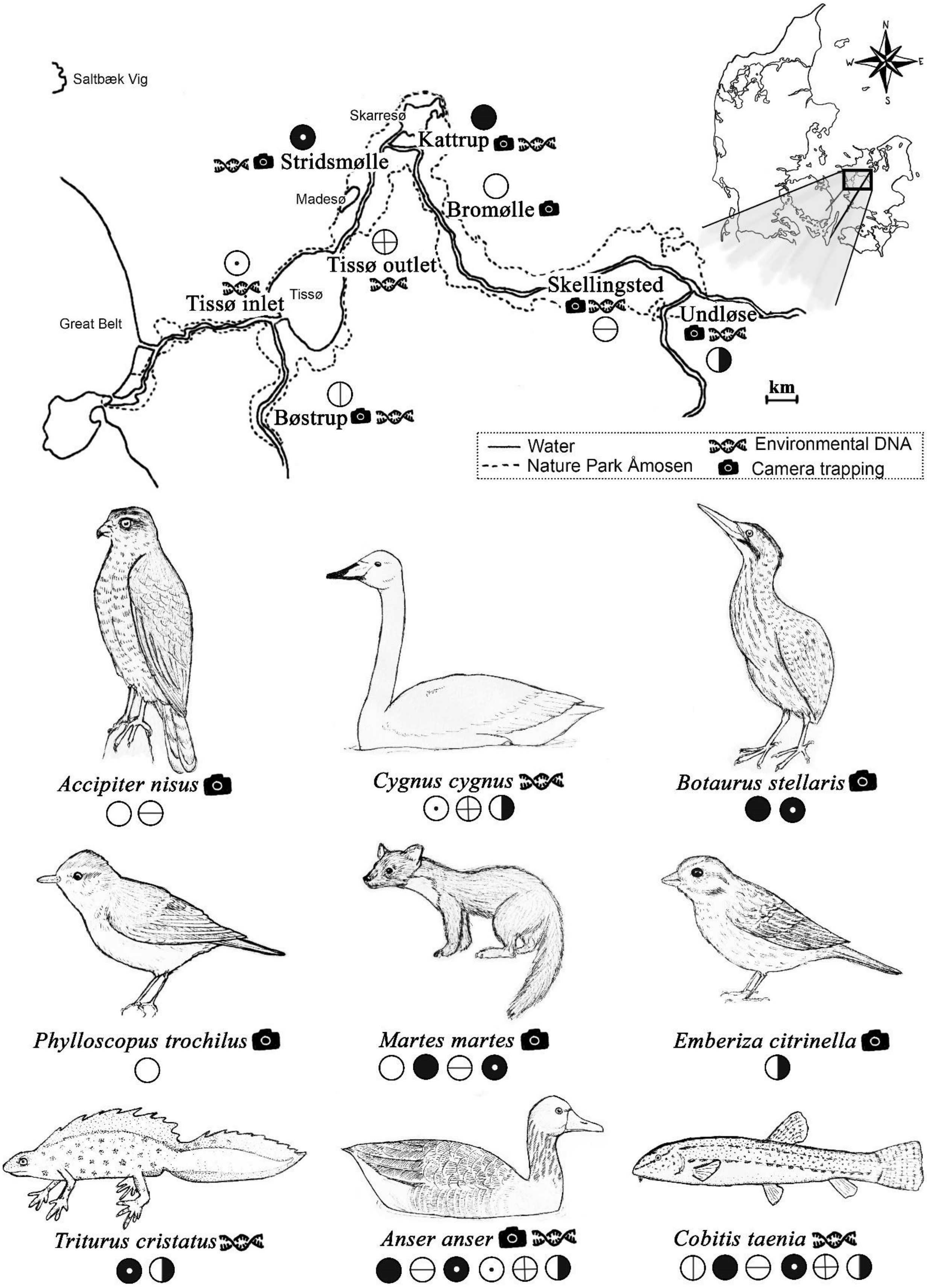
Saltbæk Vig
Skarresø
Kattrup
Stridsmølle
Madesø
Bromølle
Tissø outlet
Tissø
Tissø inlet
Skellingsted
Undløse
Great Belt
Bøstrup
km
Water
Nature Park Åmosen
Environmental DNA
Camera trapping
N
W
E
S
Accipiter nisus
Cygnus cygnus
Botaurus stellaris
Phylloscopus trochilus
Martes martes
Emberiza citrinella
Triturus cristatus
Anser anser
Cobitis taenia

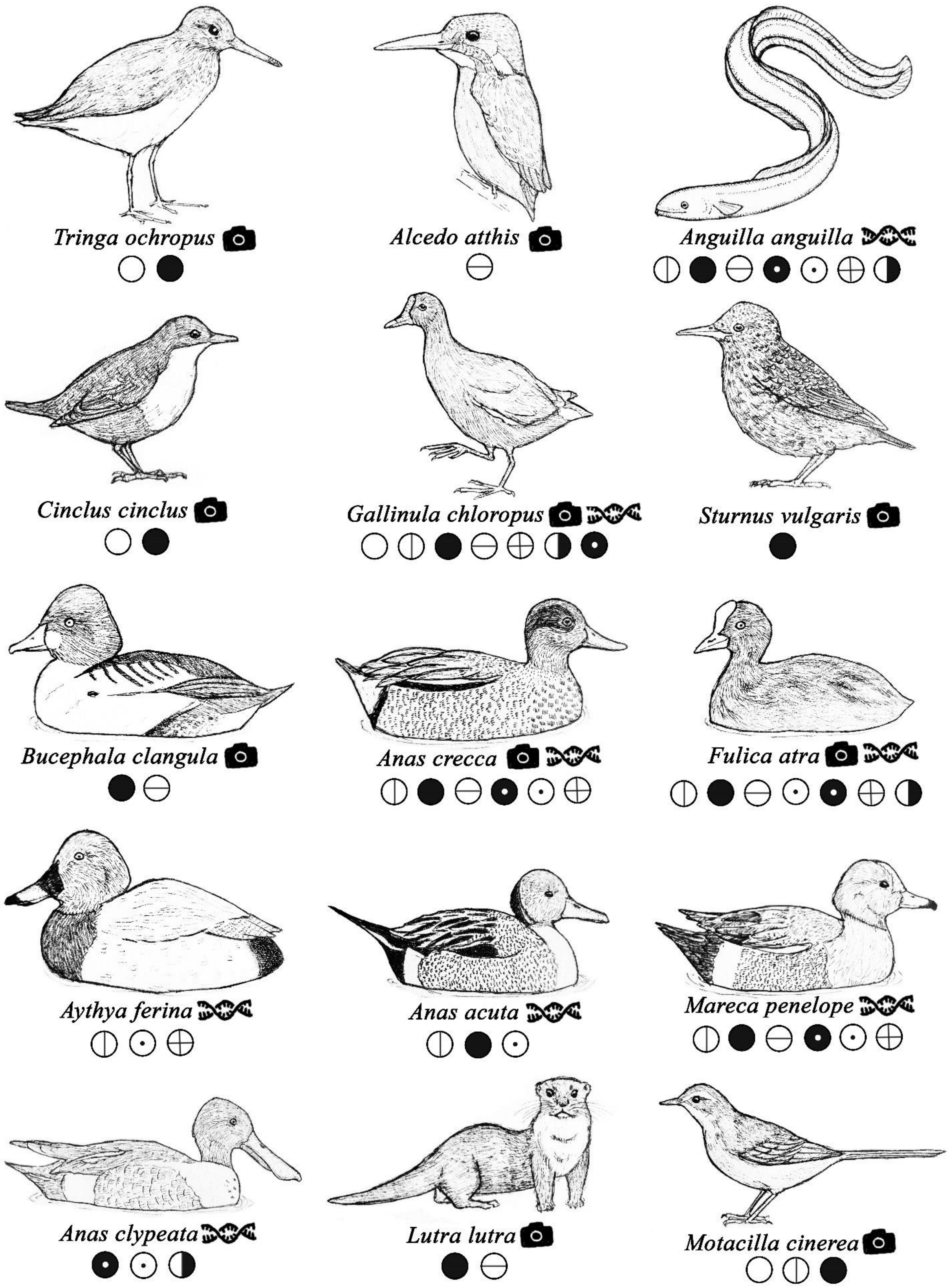

**Suppl. Figure S7** The 24 different species protected by Natura 2000 (European Commission, 2009; Naturstyrelsen, 2012) and/or on the Danish Red List (Moeslund *et al.*, 2019) detected with the use of camera trapping and/or eDNA and at which of the eight study sites. Nineteen of the species are deemed vulnerable, endangered, or critically endangered in Denmark. Furthermore, seven species are extra special for Natura Park Åmosen and have deemed the area Natura 2000 protected. The study sites are shown with circle symbols and the method of detection is indicated by either a camera or a DNA strain, representing camera trapping and eDNA, respectively. All Illustrations by AMRH. Animals are not to scale.
